# Single-nucleus multiomic analysis of Beckwith-Wiedemann syndrome liver reveals PPARA signaling enrichment and metabolic dysfunction

**DOI:** 10.1101/2024.06.14.599077

**Authors:** Snehal Nirgude, Elisia D. Tichy, Zhengfeng Liu, Rose D. Pradieu, Mariah Byrne, Luis Gil De Gomez, Brandon Mamou, Kathrin M. Bernt, Wenli Yang, Suzanne MacFarland, Michael Xie, Jennifer M. Kalish

## Abstract

Beckwith-Wiedemann Syndrome (BWS) is an epigenetic overgrowth syndrome caused by methylation changes in the human 11p15 chromosomal locus. Patients with BWS exhibit tissue overgrowth, as well as an increased risk of childhood neoplasms in the liver and kidney. To understand the impact of these 11p15 changes, specifically in the liver, we performed single-nucleus RNA sequencing (snRNA-seq) and single-nucleus assay for transposase-accessible chromatin with sequencing (snATAC-seq) to generate paired, cell-type-specific transcriptional and chromatin accessibility profiles of both BWS-liver and nonBWS-liver nontumorous tissue. Our integrated RNA+ATACseq multiomic approach uncovered hepatocyte-specific enrichment and activation of the peroxisome proliferator-activated receptor α (PPARA) – a liver metabolic regulator. To confirm our findings, we utilized a BWS-induced pluripotent stem cell (iPSC) model, where cells were differentiated into hepatocytes. Our data demonstrates the dysregulation of lipid metabolism in BWS-liver, which coincided with observed upregulation of PPARA during hepatocyte differentiation. BWS liver cells exhibited decreased neutral lipids and increased fatty acid β-oxidation, relative to controls. We also observed increased reactive oxygen species (ROS) byproducts in the form of peroxidated lipids in BWS hepatocytes, which coincided with increased oxidative DNA damage. This study proposes a putative mechanism for overgrowth and cancer predisposition in BWS liver due to perturbed metabolism.

## Introduction

Beckwith–Wiedemann syndrome (BWS) is a tissue overgrowth disorder resulting from epigenetic alterations on human chromosome 11p15^1, 2, 3^. Within the 11p15 region, there are two imprinting centers (IC1 and IC2), whose differential methylation of maternal and paternal DNA origins regulate the expression of key growth regulators insulin-like growth factor 2 (*IGF2*) and cyclin dependent kinase inhibitor 1C (*CDKN1C*). Gain of methylation at IC1 (IC1 GOM) leads to biallelic *IGF2* expression, loss of methylation at IC2 (IC2 LOM) leads to decreased *CDKN1C* expression and paternal uniparental isodisomy of chromosome 11p15 (pUPD11) leads to both increased *IGF2* expression and decreased *CDKN1C* expression^4^. Both IC2 LOM and pUPD11 cause a wide range of fetal and neonatal overgrowth of the liver, including hepatomegaly, mesenchymal hamartoma and increased incidence of hepatoblastomas^4, 5^. However, the connections between 11p15 chromosome alterations and the downstream molecular cues leading to BWS liver phenotypes have not been established.

A previous study from our lab, using bulk liver tissue, found cancer predisposition and oncologic signatures in BWS liver and hepatoblastoma^4^. This study also highlighted that the methylation profiles of liver at 11p15 can be different than the methylation profile of blood in the same patient^2,4^. This finding implies that BWS liver has altered tissue-specific molecular signatures due to 11p15-driven changes^4^. However, the molecular impact of 11p15 changes on BWS liver, are not well understood. Assessing BWS and nonBWS liver tissue in this study provided insight into the functional consequences of 11p15 alterations between the normal and the overgrown liver that contribute to the BWS-specific cancer predisposition signatures.

The liver is the largest gland and performs both endocrine and exocrine functions. In addition, it executes essential functions, including glycogen storage, detoxification/xenobiotic metabolism, cholesterol synthesis and transport, urea metabolism and secretion of plasma proteins^6, 7^. Most importantly, the liver is the key metabolic regulator that systemically governs energy metabolism^8^. About 78% of liver volume is composed of hepatocytes, the parenchymal cell type of liver. These hepatocytes coordinate their functions with non-parenchymal cell types, including cholangiocytes, endothelial cells, Kupffer cells, immune cells and hepatic stellate cells^6^. The cellular composition and intracellular signaling of liver can be comprehensively studied using single cell sequencing approaches. A number of studies have reported detailed single cell maps of adult human liver^9, 10, 11, 12^, while a study by Wesley *et al.* described different stages of liver development including hepatoblast, fetal and adult stages^11^. In the current study, we query the cellular composition and functionality/signaling of normal, nontumorous BWS liver compared to nonBWS liver, as a first step in understanding the mechanisms of BWS overgrowth in the liver. We previously reported that patients with BWS exhibit elevated alpha fetoprotein (AFP) levels in the first year of life, suggesting that BWS livers are more metabolically active^14^. This observation formed the basis and rationale to build a reference map for BWS pediatric liver at single-nuclei resolution.

We used the recent technology of multimodal single-cell sequencing, to perform parallel RNA-seq+ATAC-seq on isolated single-nuclei from flash frozen liver samples. snATAC-seq adds additional chromatin-correlated information that would otherwise be unavailable by snRNA-seq, specifically to aid in prediction of cell-type-specific cis-regulatory DNA interactions, transcription factor activity and identification of distant regulatory regions that influence gene transcription^15^. Using snRNA-seq coupled with snATAC-seq, we performed sequencing on seven human liver samples belonging to BWS and nonBWS tumor-adjacent samples, with the goal to understand the molecular signatures that define BWS liver signaling. snRNA-Seq identified similar cellular types between both groups; however, cellular pathways related to fatty acid and lipid metabolism as well as PPARA signaling were enriched in BWS liver when compared to nonBWS-liver. snATAC-seq uncovered differential chromatin accessibility for these signaling pathways, with increased accessibility in the BWS livers. This study provides a detailed map of pediatric liver composed of different cell types and explores gene expression changes in BWS liver that may drive its metabolic activity. Since BWS modelling in the mouse has limitations^16^, we utilized a BWS-iPSC model to validate our metabolic pathway patient data. Specifically, we subjected BWS-iPSC and control lines to hepatocyte differentiation, which showed the increased enrichment of fatty acid and lipid metabolism pathways in BWS lines. Our RNA-seq and real-time qPCR data demonstrated enrichment of PPARA in BWS lines relative to control lines, as differentiation proceeded. We additionally scored iPSC-hepatocytes for metabolic properties and found decreased neutral lipid droplets, concomitant with increased fatty acid β-oxidation, in the BWS lines, indicating alterations in the metabolic activity of BWS-liver. Finally, we examined proliferation and reactive oxygen species (ROS) signaling and found upregulation of proliferation, as well as an increased ROS response, along with an increase in oxidative DNA damage in the BWS hepatocytes. We propose here a possible mechanism that defines the molecular setup of neoplastic transition in BWS-livers driven by metabolic perturbations.

## Results

### Clinical cohort

The patients with BWS ranged in age from 2 months to 14 months and included samples from both males (n=2) and females (n=2). In the nonBWS cohort, patients ranged in age from one month to 8 years and included both male (n=2) and female (n=1). This study utilized cohorts with as closely matched ages as was feasible and available within the institutional biobank. Histologic review revealed similar liver morphologies between both BWS and nonBWS groups (**Supplementary Fig. 1 a, 1b**). We also calculated hepatocyte sizes in both cohorts and found that BWS liver had smaller and more numerous hepatocytes, relative to nonBWS livers (**Supplementary Fig. 1c**), which may contribute to reported BWS hepatomegaly. We confirmed the methylation status of the two imprinted regions on chromosome 11p15 (H19-IC1 and KvDMR-IC2) in nonBWS liver samples by performing pyrosequencing for KvDMR10 and H19 (**Supplementary Fig. 2a, 2b**). The nonBWS livers showed normal methylation at both IC1 and IC2. Clinical testing for BWS molecular characterization was performed in blood and non-tumor liver at the University of Pennsylvania Genetic Diagnostic Laboratory, as previously described ^17^.

### The landscape of cells in the BWS and nonBWS livers

snRNA-seq and snATAC-seq were performed on the four BWS livers and three nonBWS livers (**Fig. 1a**). All the samples were sequenced in a single sequencing run to avoid batch effect. We used Seurat (version 4)^18, 19, 20, 21^ and Harmony ^22^ packages to determine the cellular composition of the snRNA-seq samples, to annotate cells based on their transcriptional profiles, and to inform the snATAC-seq analysis. After quality control filtering and removal of putative doublets, we obtained the snRNA-seq profile for 74,315 nuclei. Using an unsupervised clustering approach, we identified 7 cell populations represented by 18 cell clusters that identified both parenchymal and non-parenchymal liver cells in both BWS and nonBWS livers (**Fig. 1b, 1c**). Their composition included hepatocytes, cholangiocytes, hepatic stellate cells, Kupffer cells, endothelial cells, and immune cells. We performed annotation using SingleR, ScType and manual annotation methods. For manual annotation, we obtained markers using the FindClusters function from Seurat (**Supplementary File 1 – Top 50 cell markers in each cluster**) and we observed cell-type-specific expression of a number of genes (**Fig. 1d, 1e**). We identified hepatocytes using known markers defined in previous liver studies, including: *ALB, ACSL4, FGG, FGA, CYP3A4, PCK1, CPS1, CP, ASGR1, APOC1* and *CYP2E1* ^9^. Using these genes, we identified seven clusters (Cluster 0, 1, 7, 17, 14, 2, 15) representing hepatocytes. Using *KRT7, KRT19* and *CDH1* markers we identified two clusters (Cluster 3, 10) representing cholangiocytes ^10^. Two clusters (Cluster 5, 8) of hepatic stellate cells were identified by *IGFBP7, PDGFRA, COL3A1* and *DCN* markers.

**Fig. 1:**
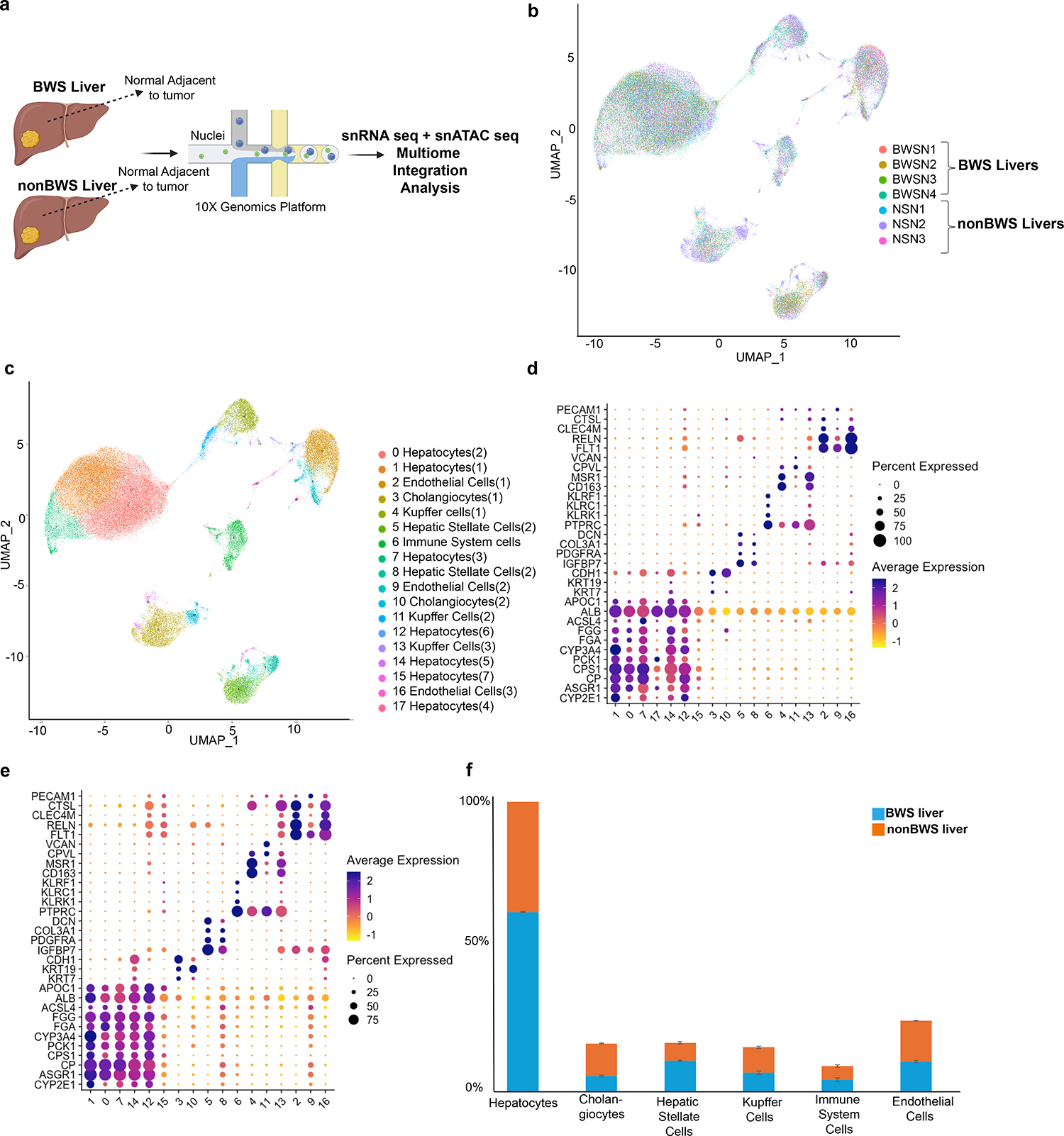
snRNA-seq profiles of BWS (n=4) and nonBWS (n=3) livers. **(a)** Schematic of our multiomics study. Single nuclei were isolated from 4 BWS and 3 nonBWS livers and prepared for snRNA-seq using the 10X Genomics technology. (**b)** UMAP visualization 74,315 liver nuclei isolated from 7 liver samples. (**c)** The uniform manifold approximation and projection (UMAP) of BWS (n=4) and nonBWS (n=3) livers using snRNA-seq demonstrates that both liver groups contain the same cell types clustered into 18 cell subpopulations. This transcriptome atlas identified major liver cell types including hepatocytes, cholangiocytes, hepatic stellate cells, Kupffer cells, endothelial cells and immune system cells in both BWS and nonBWS livers. Dot plots of snRNA-seq dataset showing gene expression patterns of cluster-enriched markers for BWS liver **(d)** and nonBWS livers. **(e).** The diameter of the dot corresponds to the proportion of cells expressing the indicated gene and the density of the plot corresponds to average expression relative to all cell types. **(f)** Percentage of cell type distribution between BWS and nonBWS livers.

*PTPRC* and *CD163* markers identified the clusters (Clusters 6, 4, 11, 13) for Kupffer cells and the immune cells^10^. Three clusters (Clusters 2, 9, 16) representing endothelial cells were identified using markers *FLT1, RELN, CLEC4M, CTSL* and *PECAM1*^10^. All cell types were identified in both liver groups (BWS and nonBWS) (**Fig. 1f**). Specifically, in BWS livers, hepatocytes represented ∼62% of total analyzed nuclei, whereas cholangiocytes, hepatic stellate cells, Kupffer cells, immune cells and endothelial cells represented 5%, 10%, 6%, 4% and 10% respectively. In nonBWS livers, hepatocytes represented ∼55% of total analyzed nuclei, whereas cholangiocytes, hepatic stellate cells, Kupffer cells, immune cells and endothelial cells represented 11%, 6%, 8%, 4% and 14%, respectively.

### Liver zonation and characterization of pediatric liver

The liver is a central metabolic organ composed of hepatocytes that exhibits a spatial distribution of functions based on oxygen gradient, a phenomenon termed as “zonation”^23^. Zonation is implicated in initiation and progression of various liver diseases^23^, and also can explain zonated damage in liver pathology^24^. It has previously been shown that hepatocytes with 11p15 alterations have changes in the liver zonation architecture^13^, hence we queried the effects of BWS alterations on hepatocyte zonation in our patient cohorts. Using human liver zonation markers previously established^9^ ^12^, a pediatric liver zonation study was performed. We classified all seven hepatocyte clusters for both liver cohorts into liver zones. Pericentral hepatocytes were mainly represented by cluster 1, based on markers *CYP2E1, SLC1A2, CYP3A4, BHMT, ABCB4,* and *GPHN* (**Fig. 2a**, **2b** and **Supplementary File 1)**. Periportal hepatocytes were mainly represented by cluster 0, 7, 14 and 17, based on the markers *HAL, SDS, C3, FGG, CRP, LEPR, CYP3A7, FGA,* and *CYP2A7* (**Fig. 2c, 2d** and **Supplementary File 1**). Cluster 15 has the smallest library size of the hepatocyte clusters (309 nuclei from both liver cohorts). The marker profile for cluster 15 did not represent any of the zonation markers and future work will be needed to clarify the origin of this cluster. Cluster 12 was a mixture of periportal and pericentral hepatocytes. We further utilized the documented functional studies for liver zones reported by Payen *et al.*^12^, to confirm the marked liver zones in both liver cohorts. The pericentral hepatocytes are mainly involved in fatty acid biosynthesis, (**Fig. 2e**), xenobiotic metabolism (**Fig. 2f**) and retinoid metabolism (**Fig. 2g**). The periportal hepatocytes are mainly involved in the secretion of plasma proteins (**Fig. 2h**). In summary, hepatocytes in BWS livers highly expressed genes involved in fatty acid biosynthesis and plasma protein secretion compared to the hepatocytes from nonBWS liver. Liver cell types were similarly represented in BWS and normal livers, but the zonation and metabolic profiles differed, suggesting altered metabolic functions in BWS livers.

**Fig. 2:**
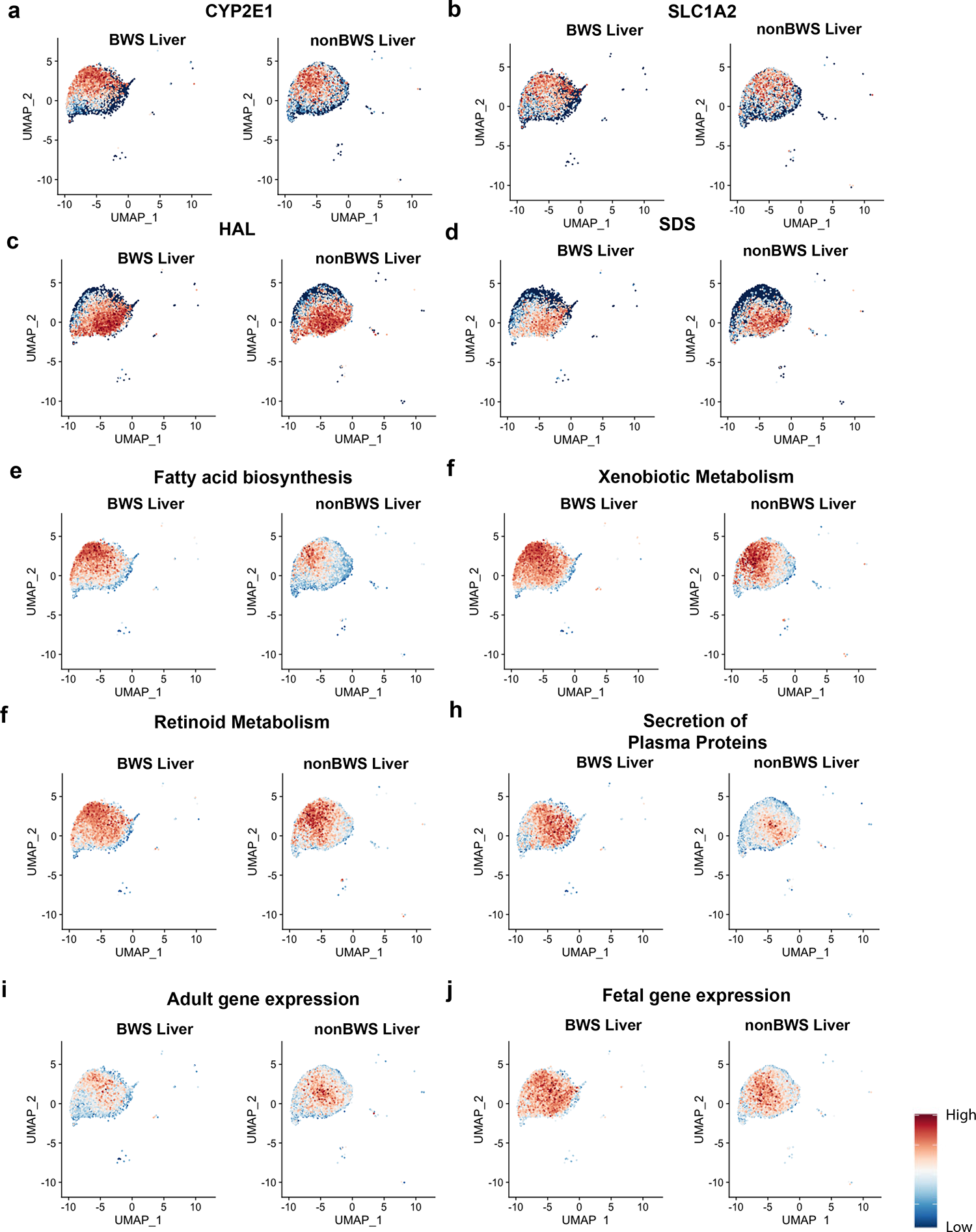
Liver zonation and developmental profile of BWS (n=4) and nonBWS (n=3) liver. Feature plot for CYP2E1 **(a)** and SLC1A2 **(b)** represent the pericentral markers whereas HAL **(c)** and SDS **(d)** represent the periportal markers. Distribution of the expression of genes associated with pericentral functions of hepatocytes like fatty acid biosynthesis **(e)**, xenobiotic metabolism **(f)**, retinoid metabolism **(g)** and periportal function of secretion **(h)** in hepatocytes. Distribution of the expression of adultl genes **(i)** and fetal genes **(j)** in BWS and nonBWS livers.

Since pediatric livers are in developmental transition between fetal and adult livers, and the developmental state of BWS livers may be abnormal^14, 25^, we wanted to investigate the expression patterns of fetal and adult liver gene signatures in our cohorts. We used the gene sets defining the fetal and adult stage of hepatocyte differentiation as described^11^. Then we generated a Feature plot representing expression of genes associated with hepatocyte differentiation. We found that both BWS and nonBWS livers expressed adult liver markers (**Fig. 2i**). However, BWS livers displayed enrichment of fetal gene signatures (**Fig. 2j**), when compared to nonBWS livers. These results imply that the developmental state of BWS livers is at an earlier stage, compared to nonBWS pediatric livers.

### snATAC-seq cell annotations are validated based on snRNA-seq data

To understand the transcriptomic regulation of genes, we performed snATAC-seq and captured the chromatin accessibility profile of individual liver cells. Since less is known about the cell type-specific chromatin accessibility profiles of liver cells, we used our annotated snRNA-seq dataset to predict the snATAC-seq cell types. Using shared barcodes from joint multiomics profiling and ArchR^26^, we performed the snATAC-seq cell annotation, which represented the 18 cell subpopulations in the snATAC-seq dataset for both liver cohorts (**Fig. 3a, 3b**). We found 25,472 snATAC-seq marker peaks enriched in cell-type specific manner in BWS liver (**Fig. 3c**) and 19,772 snATAC-seq marker peaks enriched in nonBWS liver (**Fig. 3d**). These peaks were enriched in promoter, distal, intronic and exonic regions for both livers. We then generated a heatmap of the chromatin accessibility profile of the liver cohorts (**Fig. 3e, 3f**) which revealed that the cell types within each cohort had a unique chromatin accessibility profile implying differences in their gene expression and epigenetic state.

**Fig. 3:**
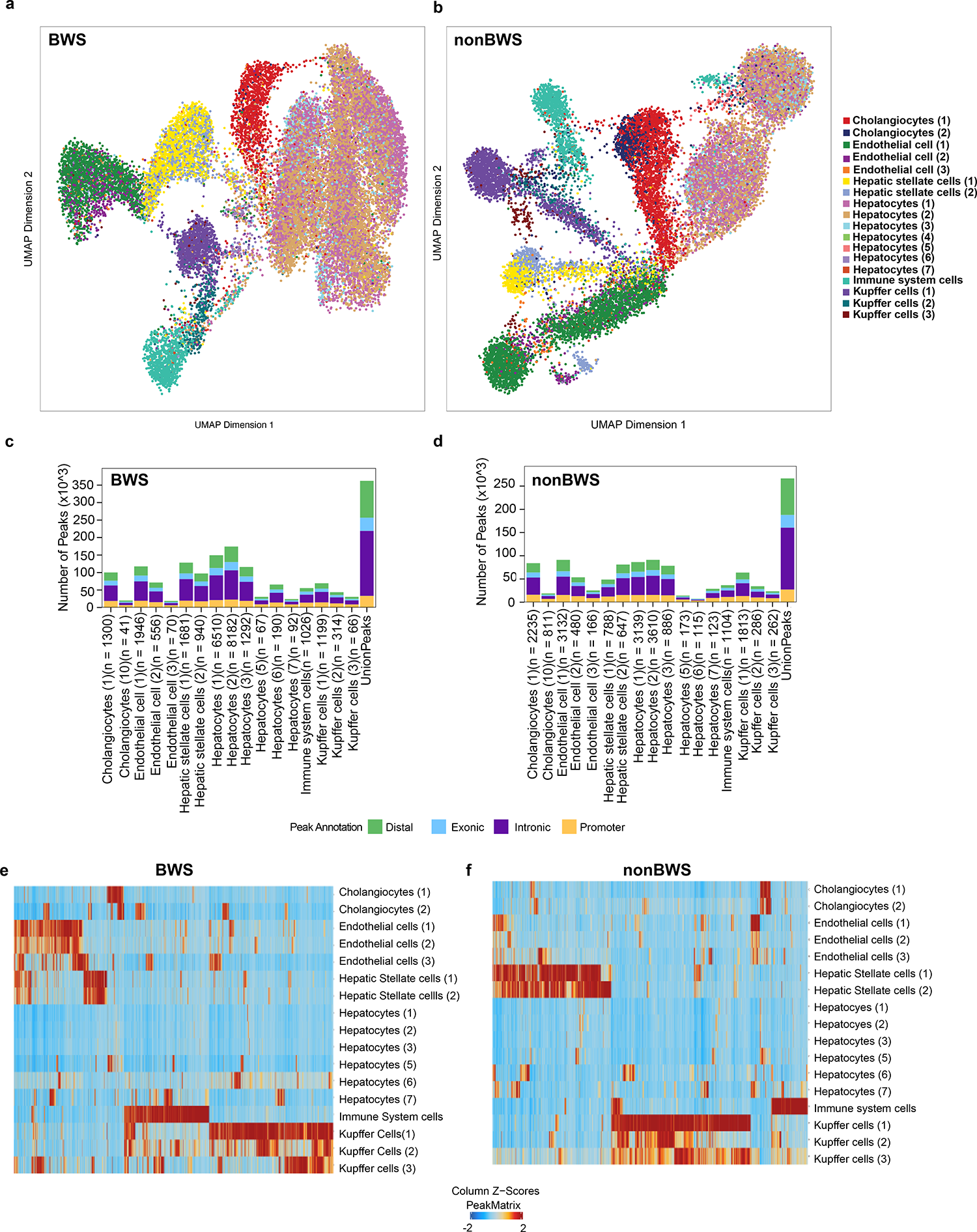
Chromatin accessibility profiles from joint snRNA-seq and snATAC-seq of the BWS(n=4) and nonBWS (n=3) livers. The uniform manifold approximation and projection (UMAP) of BWS (n=4) and nonBWS (n=3) livers using snATAC-seq with cell annotation predicted from snRNA-seq dataset for BWS liver **(a)** and nonBWS liver **(b)**. All 17 cell subpopulations representing major cell types in liver are represented. Bar plot of annotated differentially accessible region (DAR) locations for each cell type in BWS liver **(c)** and nonBWS liver **(d)**. Heatmap of average number of Tn5 cut sites and snATAC marker peaks enriched within a DAR for each cell type (left) in BWS liver **(e)** and nonBWS liver **(f).** Each column represents a marker peak.

### SCENIC plus and Gene Ontology (GO) analysis reveals altered metabolic pathways in BWS livers

To understand the underlying transcription factor – gene regulation network that makes BWS-livers distinct from nonBWS ones, we utilized single-cell regulatory network inference and clustering (SCENIC+) to construct the regulatory network from our single-nuclei multiomic dataset. We performed this analysis by subsetting the three major hepatocyte clusters (Cluster 0, 1, and 7) presented in both cohorts. By comparing the hepatocytes from both BWS and nonBWS liver groups, we identified 27 activating enhancer-regulons networks (eRegulons). (**Fig. 4a, Supplementary File 2 – List of all eRegulons regulated in hepatocytes of BWS liver**). The most significant upregulated activating eRegulons in BWS were peroxisome proliferator-activated receptor α (PPARA), its binding partner, Retinoid X Receptor Alpha (RXRA), and T cell factor/lymphoid enhancer factor (TCF7L1). The interaction network among PPARA and RXRA eRegulons is shown in **Supplementary Fig. 3**. PPARA and RXRA are involved in a variety of liver functions, including the production and breakdown of lipids and amino acids^27^, and dysfunctional PPARA signaling is implicated in hepatocyte proliferation^28^ and hepatomegaly^29^, while TCF7L1 is a WNT signaling regulator and is implicated in cell stemness^30, 31^.

**Fig. 4:**
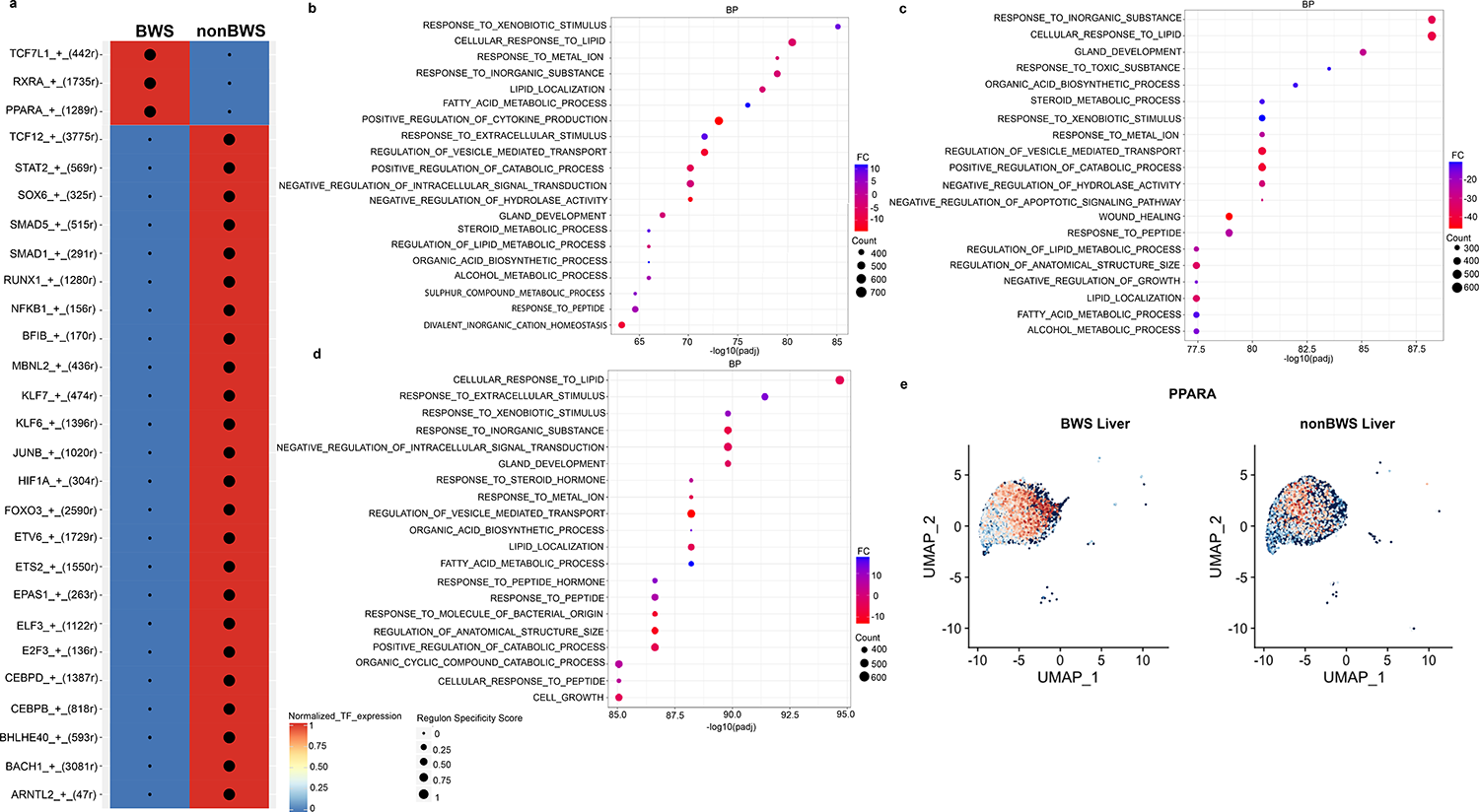
SCENIC analysis and GO term analysis for BWS (n=4) and nonBWS (n=3) livers. **(a)** Heatmap/dot plot showing transcription factor expression and group specificity of the eRegulon are displayed using SCENIC plus. Here, only activating eRegulons with calculated region-to-gene correlation coefficients (rho) bigger than 0.6 are included in the graph **(b-d)** GO terms enriched in the three hepatocytes clusters (Cluster 0, 1, 7) of BWS livers when compared to nonBWS livers. **(e)** Feature plot depicting PPARA expression in BWS and nonBWS livers.

We also performed Gene ontology (GO) term analysis using the snRNA-seq data on the cell clusters captured in the BWS livers in comparison to nonBWS livers. **Fig. 4b-4d** depicts the GO terms enriched in the Cluster 0, 1 and 7 representing Hepatocytes (2), Hepatocytes (1) and Hepatocytes (3) cell types. The GO terms enriched for rest of the hepatocyte cell clusters are provided in the **Supplementary Fig. 4**. All cell clusters representing hepatocytes in BWS livers were involved in fatty acid and lipid metabolic processes. It is well known that PPARA is a major regulator of fatty acid metabolism and lipid metabolism in liver^32, 33^. Similar to our previous data, we confirmed the overexpression PPARA in the three major hepatocytes clusters (Cluster 0, 1 and 7) in the BWS livers when compared to nonBWS livers (Fig. 4e), using the snRNA-seq dataset. Thus, SCENIC+ and GO term enrichment analysis demonstrated an involvement of PPARA signaling in BWS-liver metabolism. These findings suggest that PPARA signaling changes are a major contributor to liver alterations in BWS.

### BWS-iPSC hepatocyte model depicts the metabolic nature BWS livers

We next performed *in vitro* studies to validate our findings of PPARA dysregulation in BWS hepatocytes. We generated two BWS-iPSC lines from a parental control line using a CRISPR-Cas9 approach. All iPSC lines – BWS and control, had a normal karyotype (data not shown). We differentiated these lines down a hepatic lineage and queried the methylation status of chromosome 11p15 by pyrosequencing on Day 0 (iPSC stage), Day 10 (hepatic progenitor stage), as well as on Days 15, 18, and 20 (hepatocyte stages) (**Supplementary Fig. 5**), to ensure hepatocyte differentiation did not affect the BWS critical region. We found that our engineered BWS iPSC lines had the expected 11p15 methylation profile, which was retained during hepatocyte differentiation. We also confirmed the onset of differentiation into endoderm by performing quantitative RT-PCR at Day 5 of differentiation for relevant markers (*CXCR4, FOXA2,* SOX17) (**Supplementary Fig. 6**). We further examined the efficiency of hepatocyte differentiation by querying the expression profile of several functional hepatocyte genes on days 15 and 20 of differentiation by performing RNA-seq (**Supplementary Fig. 7**). We observed comparable expression of the markers between control and BWS lines, demonstrating no defects in early hepatocyte differentiation programming exist in BWS. Then we performed GO term enrichment analysis on Day 15 and Day 20 hepatocyte-differentiated lines. Compared to the control line on Day 15 and Day 20, we found activation of fatty acid and lipid metabolic processes (**Fig. 5a-d**) in line with GO term enrichment analysis on BWS-liver snRNA-seq analysis. Additionally, we monitored the changes in cellular morphology and cell size during the hepatocyte stages of differentiation between control and BWS lines (**Supplementary Fig. 8**). Cellular morphology imaging revealed possible changes in the lipid droplet accumulation, while size analysis displayed smaller hepatocytes in BWS-altered cells (**Supplementary Fig. 8**), similar to our observations in BWS patient livers (**Supplementary Fig. 1**). Since PPARA was enriched in our multiomic study, and is major regulator of lipid metabolism in liver, we studied expression of PPARA in our RNAseq data and also performed qRT-PCR for PPARA on Day 10, Day 15, Day 18 and Day 20 of hepatocyte differentiation (**Supplementary Fig. 9)**. We found that PPARA expression increased over the course of differentiation in BWS cells compared to control (**Fig. 5a-d**), consistent with our multiomic analyses on patient livers.

**Fig. 5:**
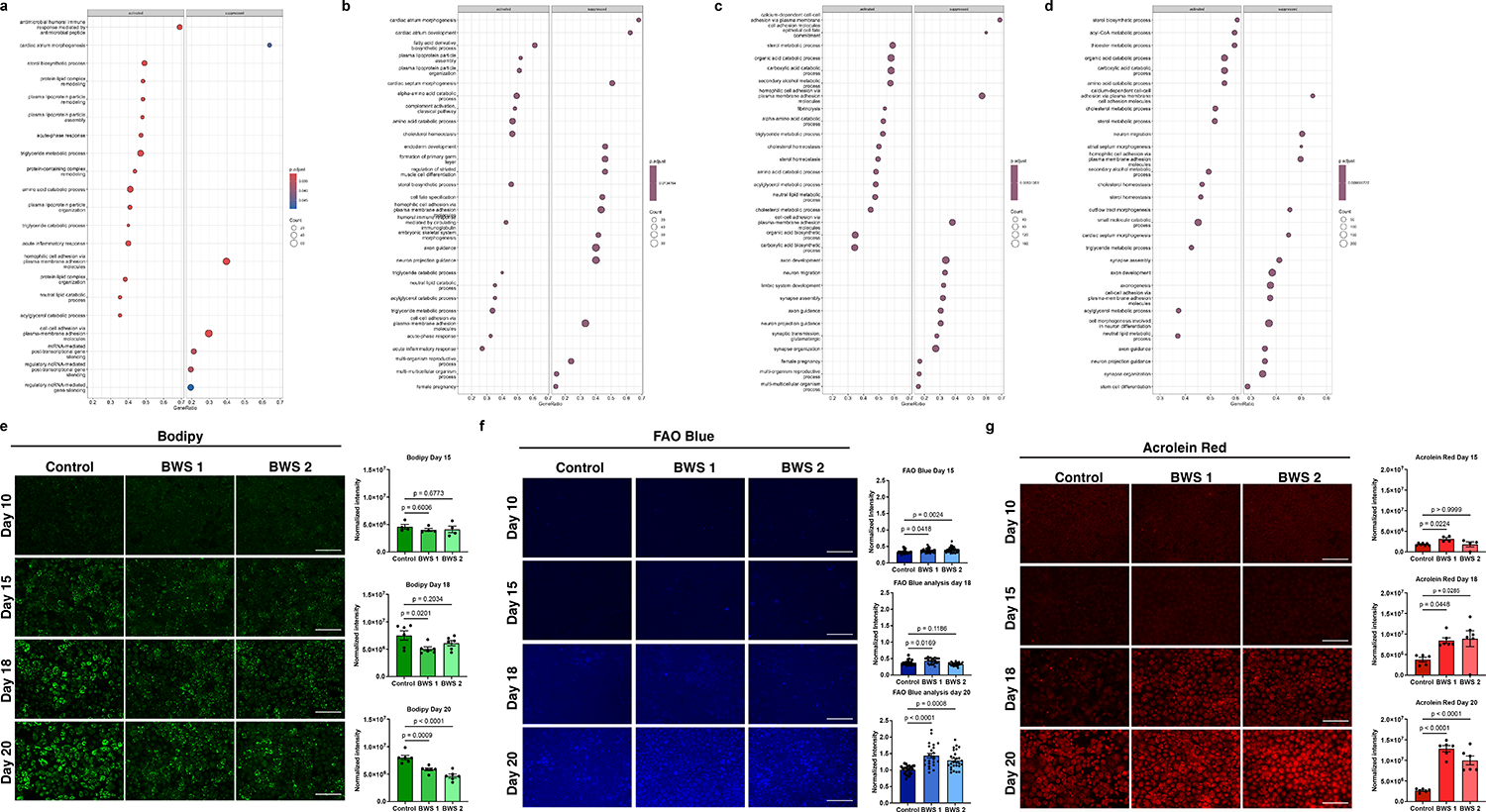
Hepatocyte-specific BWS effects on PPARA and fatty acids *in vitro*. GO term enrichment plot for Day 15 of hepatocyte differentiation in BWS 2 (**a**) and BWS 1 (**b**) line when compared to Day 15 of hepatocyte differentiation in the control line. GO term enrichment plot for Day 20 of hepatocyte differentiation in BWS 2 (**c**) and BWS 1 (**d**) line when compared to Day 20 of hepatocyte differentiation in the parental line. (**e-g**) Live cell imaging of control, BWS 1, and BWS 2 cells undergoing differentiation and stained with Bodipy (**e**), FAO Blue (**f**), and Acrolein Red (**g**). Quantification for days 15, 18, and 20 of differentiation are shown beside the image and represent mean ± SEM. p values were generated by one way ANOVA.

### BWS iPSC-derived hepatocytes exhibit altered lipid metabolism

Since PPARA is involved in cellular processes that include lipid metabolism^32, 33^, we next examined the effects of BWS-specific PPARA increases on several lipid processes. We first confirmed our morphological study that suggested differences in lipid droplet accumulation in BWS hepatocytes. Bodipy 493, a cell permeable nonpolar dye, accumulates in neutral lipid droplets and fluoresces when bound to neutral lipids^34, 35^. We found that the fluorescence intensity following Bodipy staining increased over time in control and BWS lines (**Fig. 5e**) indicating increases in lipid droplet formation. However, this increase was much greater in control lines when compared to BWS lines. This finding can be explained by an increase in fatty acid breakdown (β-oxidation) and/or less production of fats. Since PPARA targets include enzymes in the β-oxidation pathway, we assessed lipid breakdown by utilizing a fluorescent indicator of this process, FAO blue ^36, 37^. Staining cells with this dye revealed that β-oxidation increases in both control and BWS hepatocytes over our differentiation time course (**Fig. 5f**). By Day 20, BWS hepatocytes exhibited more β-oxidation when compared to control hepatocytes, which exhibits the highest *PPARA* expression begins increasing in these cells (**Supplementary Fig. 9**). These findings are in agreement with our omics data that displayed increased lipid metabolism gene enrichment in BWS liver. We also observed upregulated expression of genes related to fatty acid β-oxidation, including ACOX1, ACOX2, ACSL1, ACSL4, XBP1, CAT and EHHADH on Days 15 and 20 of hepatic differentiation in BWS-iPSC lines relative to the parental control line (**Supplementary Fig. 10**). In aggregate, these findings depict an altered metabolic environment in BWS hepatocytes, with increased breakdown of lipids.

### BWS iPSC-derived hepatocytes display increased ROS signatures

A consequence of β-oxidation is the generation of endogenous reactive oxygen species (ROS)^38, 39^, which, among other effects, can lead to lipid peroxidation. To assess this possibility, we performed acrolein red staining, which uses a fluorescent indicator to measure lipid peroxidation caused by oxidative stress^40, 41^. As observed in **Fig. 5g**, BWS hepatocytes showed enrichment in acrolein red staining when compared to control hepatocytes, particularly at later time points examined. These findings are indicative of a role of increased PPARA in the BWS-hepatocytes contributing to increased fatty acid oxidation, which subsequently coincides with increase in lipid peroxidation, a marker of oxidative stress. To confirm our findings, we studied expression oxidative stress-related genes in control and BWS hepatocytes by RNAseq. We observed upregulation in the expression of ROS scavenging enzymes like GPX1, GPX3, GPX4, GPX7, SOD2, SOD3 and NUDT1 in BWS lines of Day 15, Day 20 differentiated cells when compared to the control line **Supplementary Fig. 11**). Since BWS livers are more prone to tumorigenesis, we next examined the level of oxidative DNA damage, by staining Day 20 differentiated hepatocytes with 8-oxoguanine antibody. We found BWS hepatocytes exhibited more oxidative DNA damage, relative to control, and this finding was correlated with increased expression of OGG1 in BWS lines (**Supplementary Fig. 12**). In aggregate, these findings demonstrate BWS cells are more metabolically active and exhibit more oxidative stress and DNA damage, which may contribute to the development of hepatoblastomas.

## Discussion

The purpose of this study was to characterize BWS normal liver tissue and discover BWS-specific transcriptional and epigenetic alterations compared to nonBWS livers. We examined the transcriptomic profile, in conjunction with chromatin accessibility profile of BWS and nonBWS pediatric liver in an unbiased manner using a single nuclei multiomic sequencing approach. This approach revealed cell-type-specific enrichment of the PPARA – a liver metabolic regulator^32, 42^. To confirm our findings, we generated a BWS-iPSC model and differentiated the generated iPSCs into hepatocytes. Our *in vitro* data demonstrates the dysregulation of lipid metabolism in BWS-hepatocytes, which coincided with observed upregulation of PPARA during hepatocyte differentiation. BWS liver hepatocytes exhibited decreased neutral lipids, increased fatty acid β-oxidation, and ROS signatures, relative to controls. We speculate that the altered liver metabolism is major contributor for hepatomegaly and cancer predisposition to hepatoblastoma in BWS liver.

We captured a comprehensive map of 74,315 nuclei from seven livers that included livers from both patients with and without BWS. Using the markers for liver cells described in adult liver studies ^9, 10, 12^, we identified both parenchymal and non-parenchymal liver cells in both the cohorts. Hepatocytes, the parenchymal cells, perform the liver vital liver functions based on their location and the oxygen gradient. This phenomenon is defined as “liver zonation”^24^ that informs the liver functionality^24, 43^, initiation and progression of liver disease^23^ and liver damage^24^. Non-alcoholic fatty liver disease (NAFLD), hepatic steatosis (NASH), cirrhosis, drug and alcohol-induced hepatotoxicity, as well as parasite infection-induced hepatic fibrosis all initiate in the pericentral area, while hepatocellular carcinoma (HCC) exhibits alterations in liver zonation via Wnt/β-catenin signaling^44^. Thus, liver zonation studies may be informative in identifying the underlying mechanism of disease. Based on zone specific markers validated by Halpern *et al.*^45^ in mouse liver and Payen *et al.*^12^ in adult human liver, we were able to outline the liver zone-specific hepatocytes in the BWS and nonBWS pediatric livers. We found the markers HAL and SDS clearly outlined the periportal hepatocytes while CYP2E1, SLC1A2 outlined the pericentral hepatocytes. Payen *et al.* also described genes for zone specific liver functions like xenobiotic metabolism, fatty acid biosynthesis and retinoid metabolism for pericentral hepatocytes and secretion of plasma proteins for the periportal hepatocytes^12^. We observed enrichment of these liver functional genes in both the periportal and pericentral hepatocytes in BWS livers compared to the nonBWS livers. These data suggest that BWS livers have a distinct, dysregulated metabolic program that affects both periportal and pericentral hepatocytes. However, further studies and disease models are required to decipher the role of liver zonation in BWS-hepatocytes to predict the BWS liver pathology.

To further understand the gene regulation differences, we used an integrated approach by combining the snRNA-seq and snATAC-seq data. We performed the SCENIC+ plus analysis by integrating our snRNA-seq and snATAC-seq data to understand the transcription factor/gene regulation network that defines BWS liver, where we found activation of PPARA and RXRA. PPARA is a ligand-activated nuclear receptor highly expressed in the liver parenchymal cell population i.e. the hepatocytes^33, 46^. PPARA is known to form heterodimers with RXRA and functions as a master regulator of hepatic lipid metabolism, including fatty acid synthesis, fatty acid β-oxidation, fatty acid uptake and binding, lipid droplet metabolism, lipoprotein metabolism, glycerol metabolism and cholesterol/bile metabolism, amino acid metabolism, inflammation, and energy metabolism^46, 47^. Our GO term enrichment studies in the hepatocytes comparing BWS to nonBWS liver demonstrated an enrichment of fatty acid and lipid metabolic pathways in BWS hepatocytes, indicative of PPARA involvement in the dysregulation of these processes in BWS liver. To validate our findings, we investigated PPARA functions in iPSCs differentiated into hepatocytes. Our BWS cells behaved similarly to our BWS patient samples and significantly upregulated PPARA at later stages of the hepatic differentiation process.

To understand the impact of PPARA upregulation in downstream metabolic processes in our model, we assayed the lipid accumulation, fatty acid β-oxidation and lipid peroxidation in the BWS-iPSC differentiated hepatocytes and found changes in all examined lipid metrics in BWS hepatocytes. A hallmark of hepatocyte functionality is the accumulation of neutral lipid droplets^48,49^. In our BWS iPSC hepatic differentiation system, we observed neutral lipid accumulation in hepatocytes, particularly at later time points of differentiation. However, as compared to the control line, which had progressive lipid droplet formation during differentiation, the BWS-hepatocytes exhibited reduced lipid droplet accumulation at all stages of differentiation. These findings can be explained by either a reduction in the production of lipid droplets, an increase in β-oxidation of lipids, or a combination of both. We assayed the relative level of β-oxidation during our differentiation time course. We found increased β-oxidation in BWS cells, particularly around day 20, which coincided with maximal PPARA gene expression compared to control cells. Increased fatty acid oxidation is a major source of ATP^50^ leading to increased metabolism which might contribute to hepatomegaly, a common clinical feature observed in BWS^2^. Given the critical role of the liver in regulating energy both within itself and systemically throughout the body, the regulation of insulin signaling in other BWS affected organs may tie into the defects in metabolism we observe in BWS liver like increased AFP levels in the first year of life of patients with BWS^14^.

Mitochondrial β-oxidation is a primary source of endogenous reactive oxygen species (ROS) production^38, 39^. Excessive ROS/free radicals can lead to oxidative stress in cells; however, cells can adapt by metabolic reprogramming and survive^51^. Even moderate increases in ROS can favor the induction of cell proliferation^52^, which is essential for tumor formation. Additionally, ROS can also induce DNA damage leading to mutagenesis in nucleobases like guanine and can ultimately lead to tumor-initiating events^53^. We observed increased oxidative DNA damage, as well as upregulation of the OGG1 gene (**Supplementary Fig. 12**). This gene encodes 8-oxoG DNA glycosylase 1, whose product is responsible for the excision of 8-oxoguanine, a mutagenic base byproduct which occurs as a result of exposure to ROS^54^. The combination of increased oxidative DNA damage, increased lipid peroxidation, and upregulation of ROS scavenger genes in BWS hepatocytes all agrees with increased ROS production in these cells^52, 55^. ROS, in combination with increased lipid peroxidation, has also a pathological impact in various other cancers including astrocytic and ependymal glial tumors^56^, colorectal cancer^57^, kidney tumor^58^ and breast cancers^59^. In liver, ROS and lipid peroxidation products are known to promote the differentiation and activation of hepatic stellate cells (HSCs) which is important in the transition to cancer in hepatocellular carcinoma ^60, 61^. However, the effects on ROS on the formation of hepatoblastomas are unknown. We propose that the combination of the increased proliferative state of BWS liver cells (**Supplementary Fig. 13**), in combination with the PPARA-driven metabolomic changes, results in an increased oxidative stress environment, which can damage DNA and contribute to the transition from a cancer predisposition state to hepatoblastoma formation. Further functional studies will be required to confirm this hypothesis in BWS.

One limitation of this study is small sample size and access to frozen samples. snRNA-seq has the ability to capture both parenchymal and nonparenchymal cells with high resolution and hence opens an avenue to study these cells in diseased and non-diseased state^10^. This approach has allowed us to examine our banked samples with single nuclei resolution. However, even with the sample size limitation, we were able to replicate and extend our results in an *in vitro* model of BWS liver, demonstrating the validity of our findings. In subsequent studies, we will broaden our findings using a single-cell approach on freshly isolated tissue to capture more RNA signatures and underrepresented cell types in pediatric liver. Despite these limitations, our data are the first description of the transcriptomic profile in conjunction with chromatin accessibility profile of BWS and nonBWS pediatric livers. We identified unique functional pathways enriched in the BWS liver that define its distinct metabolic function.

## Conclusion

The current study generated, to the best of our knowledge, the largest human snRNA-seq+snATAC-seq dataset currently available comparing BWS and nonBWS pediatric liver. Taken together, our transcriptional map of pediatric liver provides a framework for understanding the metabolic nature of the BWS liver when compared to nonBWS liver. Our multi-omics approach detected unique cell states within the liver and redefines cellular heterogeneity in BWS and nonBWS-livers. This study also showed that the PPARA-driven metabolic state of BWS livers could be a major contributor towards BWS liver cancer predisposition. Our findings also nominate novel prophylactic therapies, such as PPARA modulators^62^, for consideration in the prevention of BWS hepatoblastoma.

## Methods

### Patients and Samples

All the samples and clinical information from patients with BWS were collected through the BWS Registry. This registry is established at the Children’s Hospital of Philadelphia under the institutional review board protocol (IRB 13-010658). The liver samples of patients without BWS were collected under the protocol (IRB 21-018450_AM13) titled-Precision Medicine in Childhood Cancer Registry and Biorepository (PMCR). Informed consent was obtained for the collection of clinical information and the samples comprising this study. The limited clinical information about the nonBWS samples was extracted from the electronic clinical record through an honest broker. These samples were selected from individuals who did not have BWS **(Supplementary Table 1)**. BWSN1, BWSN2, BWSN3, BWSN4 represent the BWS liver cohort and NSN1, NSN2, NSN3 represent the nonsyndromic nonBWS liver cohort. The methylation profile for nonBWS cohort was confirmed normal at IC1 and IC2 by pyrosequencing (**Supplementary Fig. 2**).

The liver samples were collected from CHOP patients during their surgical hepatoblastoma resection and were confirmed as normal adjacent tissue beyond the tumor margins by a clinical pathologist. All the samples were snap-frozen in liquid nitrogen and stored at −80°C. Clinical testing for BWS molecular characterization was performed in blood and normal adjacent liver at the University of Pennsylvania Genetic Diagnostic Laboratory, as previously described ^17^.

### Single Nuclei isolation

For Multiome sequencing (snRNA-seq+snATAC-seq), nuclei were isolated with Nuclei Lysis buffer (10mM Tris-HCl, 10mM NaCl, 3mM CaCl_2_, 0.1% NP-40, 0.1% Tween-20, 1% BSA, 1mM DTT, all in 1x PBS supplemented with RNase inhibitor (1U/μL, Sigma). ∼40-45 mg tissue was homogenized using a Dounce homogenizer in 1ml of ice-cold Nuclei Lysis buffer and incubated on ice for 10 min with an additional 2mL of lysis buffer. The homogenate was filtered through a 30 μm cell strainer followed by a 100μm cell strainer. Nuclei were centrifuged at 500*xg* for 5 min at 4°C. The pellet was resuspended, washed with 1 ml of wash buffer (1x PBS +1% BSA, 1mM DTT supplemented with RNase inhibitor). The resultant nuclei suspension was FACS sorted using 7-Aminoactinomycin D (7-AAD) dye as a singlet discriminator on FACS Jazz cell sorter (BD Biosciences). 600,000 sorted events were collected in 10% BSA in 1X PBS solution supplemented with RNase inhibitor. After sorting, the nuclei pellet was centrifuged at 500*xg* for 5 min at 4°C using wash buffer and subjected to permeabilization in 100μL 0.1X permeabilization buffer (10mM Tris-HCl, 10mM NaCl, 3mM CaCl_2_, 0.1% NP-40, 0.1% Tween-20, 1% BSA, 1mM DTT, and 0.001% Digitonin, all in 1x PBS) for 2 min on ice. The suspended nuclei were centrifuged at 500*xg* for 5 min at 4°C using the wash buffer and resuspended in 40μL diluted nuclei buffer from 10X Genomics.

### Library Preparation and sequencing

10X Chromium libraries were prepared according to the manufacturer’s protocol by CHOP’s Center for Applied Genomics (CAG) core. Briefly, next-generation sequencing libraries were prepared using the 10x Genomics Chromium Single Cell Multiome Assay for Transposase-Accessible Chromatin (ATAC) and Gene Expression Reagent kit v1. ATAC libraries were uniquely indexed using the Chromium single Index Kit, pooled, and sequenced on an Illumina NovaSeq 6000 sequencer in a paired-end, dual indexing run. Gene expression libraries were uniquely indexed using the Chromium dual Index Kit, pooled, and sequenced on an Illumina NovaSeq 6000 sequencer in a paired-end, dual indexing run. Sequencing for each library targeted 20,000 mean reads per cell. Data was then processed using the Cell Ranger pipeline (10x Genomics, version 3.1.0) for demultiplexing and alignment of sequencing reads to the (reference hg38) transcriptome and creation of feature-barcoded count matrices for gene expression and ATAC data.

### Single-nuclei RNA/ATAC 10X Multiome Sequencing

#### Single-nuclei RNA and single-nuclei ATAC-seq data preprocessing and quality control

The subsequent data processing involved Seurat package (version 4.9)^18, 19, 20, 21^ in R (version 4.3). Samples with transcript counts outside the 200-50000 range, >5% mitochondrial reads, transcriptome start site (TSS) enrichment < 4, RiboRatio > 5%, and ATAC fragments < 1000 were excluded. Doublets were identified and removed using the DoubletFinder_v3 functions from the DoubletFinder R package (version 2.0.3) ^63^.

#### Single-nuclei RNAseq Analysis Processes

After performing initial quality control (QC) and removing doublets, the single nuclei RNA sequencing (scRNAseq) data was consolidated into a unified count matrix. Utilizing Seurat’s built-in functionalities, we initiated data processing. Initially, we employed “NormalizeData” to normalize the dataset. Subsequently, we applied data scaling (“ScaleData” function), accounting for mitochondrial reads (“perc.mito”) and total molecule counts (“nCount_RNA”) through regression. Next, we utilized “FindVariableFeatures” to select the top 2000 features and conducted Principal Component Analysis (PCA) using the “RunPCA” function, retaining the top 30 principal components for dimensional reduction. The resulting output served as input for Harmony ^22^ to mitigate batch effects across 30 iterations using “RunHarmony”. The harmonized data underwent cell clustering via Seurat’s functions “RunUMAP”, “FindNeighbors”, and “FindClusters”, employing the top 30 principal components at a resolution level of 0.5. For each cluster, we identified the top 50 marker genes and their corresponding expression levels. After initial QC and doublets removal, scRNAseq data were combined into a single count matrix. Seurat built-in functions were employed for data normalization, scaling (“ScaleData” function) by regressing out mitochondrial reads (“perc.mito”) and total molecule counts (“nCount_RNA”). Harmony (30 iterations) was applied to remove batch effects (“RunHarmony”). Clustering, dimensionality reduction, and marker gene identification were performed using Seurat functions: “RunUMAP”, “FindNeighbors”, and “FindClusters”. We took top 30 principal components with resolution level at 0.5. For each cluster, we reported top 50 marker genes and their expression levels. We used the AddModuleScore feature from Seurat package to study the distribution of the expression of the genes associated with a given liver function or state.

#### Single-nuclei RNAseq Cell Type Identification

We utilized two R packages to assist in defining the cell types within the obtained clusters. Firstly, SingleR (version 1.0.1) ^64^ employed the Human Primary Cell Atlas (via the R library celldex, version 1.10.1) to reference transcriptome databases, facilitating unbiased auto cell type recognition. Secondly, ScType (available at https://github.com/IanevskiAleksandr/sc-type) ^65^ provided another R package offering ultrafast automated methods for defining cell types in single-cell data. Additionally, we manually examined the marker genes from each cluster, combined with the outputs from SingleR and ScType, to assign final cell type labels to each identified cluster.

#### Single nuclei RNAseq Gene and Functional Enrichment Comparing Syndromic Patients and NonSyndromic Controls

Gene expression levels were directly compared between the Syndromic normal liver patient group (BWSN) and non-syndromic normal liver controls (NSN) within each cluster. To conduct this comparison, we utilized the Single Cell Pathway Analysis (SCPA) R package ^66^, which facilitated gene functional enrichment analysis between BWSN and NSN. Gene functional annotation relied on the Molecular Signatures Database (MSigDB) from the Broad Institute, which includes curated gene sets. Our analysis focused specifically on three collections: “HallMark”, “Canonical Pathway”, and “Gene Ontology”. Each function was ranked based on its significant adjusted p-value from the enrichment test.

#### Single-nuclei ATACseq Data Analysis

Analysis of single nuclei ATACseq data was performed using the ArchR package (version 1.0.2) (version 1.0.2) ^26^. Cell type labels obtained from scRNAseq were transferred to ATACseq based on barcodes. Initially, cells within each cell type were aggregated to generate bulk ATACseq-type data using the “addGroupCoverages” function in ArchR. Chromatin accessibility peaks were identified using the “addReproduciblePeakSet” function, utilizing the MACS2 peak identification pipeline with the human genome build hg38 as a reference. Peak properties were then added with the “addPeakMatrix” function, followed by peak calling using the “getMarkerFeatures” function, which incorporated “TSSEnrichment” and “number of fragments” as covariates, and employed the Wilcoxon test as the statistical method. Marker peaks were filtered based on an absolute log fold change (log2FC) greater than 1 and a false discovery rate (FDR) less than 0.01. Co-accessibility analysis was conducted using the “getCoAccessibility” function, with a correlation cutoff set at 0.3 and a resolution of 1. Predicted target genes for each scATAC peak were generated using the “addPeak2GeneLinks” function with Harmony output. Subsequently, the “getPeak2GeneLinks” function was applied, using a correlation cutoff of 0.3 and a resolution of 1. Enriched motifs were identified using the “addMotifAnnotations” function, utilizing the “cisbp” motif collection and gene annotations from EnsDb.Hsapiens.v86. This was followed by the “peakAnnoEnrichment” function, with a cutoff threshold set at FDR < 0.1 and Log2FC > 0.5. Finally, Footprint analysis was conducted using the “getFootprints” function.

#### Inference of gene regulatory network

To infer transcription factor activity and identify potential corresponding candidate enhancer regions and regulated genes network (eRegulons) using our single-cell multi-omic data, we utilized single-cell multi-omic inference of enhancers and gene regulatory networks (SCENIC+)^67^. To identify the differential eRegulon between BWS normal hepatocytes and non-syndromic hepatocytes, we isolated hepatocytes for the SCENIC+ analysis according to our annotated cell clusters as previously described. snATAC-seq data was first processed using pycisTopic^67^ to identify differentially accessible regions (DARs) and sets of co-accessible regions (termed topics). A model of 40 topics were selected for motif enrichment analysis using pycisTarget^67^. Motif enrichment analysis is performed using precomputed motif database and the motif-to-transcription factor annotation database from pycisTarget. eRegulons were inferred by SCENIC+ using previously processed snRNA-seq data, calculated DARs and topics, and the enriched motifs. eRegulons with negatively correlated enhancer accessibility and gene expression was filtered out. The final network was constructed using Cytoscape^68^.

#### Induced Pluripotent Stem Cell (iPSC) generation

A parental control iPSC line (Sv20) derived from an adult male peripheral blood as described^69^. This iPSC Sv20 line was genetically modified using CRISPR/Cas 9 approach to delete a portion of the second imprinting center of the maternal chromosome to create imprinted center 2 loss of methylation (IC2 LOM) lines. Two lines (3H5, 3D1, referred to as BWS 1 and BWS 2, respectively) were generated with this strategy and were confirmed by genotyping. Karyotyping was performed to confirm a normal chromosome complement. The methylation pattern at IC1 and IC2 were confirmed by pyrosequencing.

#### iPSC differentiation

Parental control line and two BWS IC2 LOM lines (BWS 1, BWS 2) were plated on laminin-coated plates. 0.5×10^6^ cells were seeded in a 24-well plate. Cells were subjected to a 20-day protocol for hepatocyte differentiation using the STEMdiff^TM^ hepatocyte kit (Stem Cell Technologies), according to the manufacturer’s instructions. Cells were imaged by brightfield/phase microscopy throughout the differentiation protocol using an EVOS M7000 imaging system to monitor differentiation.

#### RNA isolation, cDNA synthesis, and quantitative RT-PCR (qRT-PCR)

Control and BWS IC2 LOM iPSCs were plated in 12 well laminin-coated plates. Cells were differentiated for 5 days with the STEMDiff hepatocyte kit (Stem Cell Technologies), according to the manufacturer’s instructions. RNA was isolated with Trizol (Thermo Fisher Scientific) and reverse transcribed to cDNA with an iScript cDNA synthesis kit (Bio-Rad). Quantitative real time PCR was conducted in technical triplicates or quadruplicates and 4 biological replicates for each data point. Gene expression was quantified using a multiplexed ddCT method, using *B2M* as a housekeeping gene. All primers with TaqMan probes were purchased from Thermo Fisher Scientific. Gene assay ID’s: FAM-*CXCR4* (Hs00976734_m1), FAM-*FOXA2* (Hs00232764_m1), FAM-*SOX17* (Hs00751752_s1), VIC-*B2M* (Hs00187842_m1) and FAM-*PPARA* (Hs00947536_m1).

#### Live cell staining, imaging, and analysis

iPSCs were differentiated and used for assays at timepoints of Day 10, Day 15, Day 18, and Day 20 of the hepatocyte differentiation protocol. Cells were stained for at 37°C in the dark with either 2.5 μM Bodipy 493 (to assess neutral lipids; Invitrogen; one hour), 2 μM FAOBlue (to assess fatty acid Beta oxidation; Diagnocine; two hours) or 10 μM Acrolein red (lipid peroxidation and oxidative stress; Diagnocine; one hour) in Hepes-buffered saline (HBS). Cells were washed with HBS and imaged with an EVOS M7000 imaging system equipped with a GFP filter, a BFP-tag filter, and a Texas red filter, respectively. Data was collected by imaging of 4 random fields from at least 2 biological replicates per time point. Analysis of intensities were conducted in FIJI and normalized to cell number by independent investigators.

#### Bulk RNAseq and analysis

Undifferentiated iPSCs (Sv20, 3D1, 3H5) and iPSCs (Sv20, 3D1, 3H5) undergoing hepatocyte differentiation at D15 and D20 were collected in two biological replicates from 24-well plates. Briefly, cells were washed with cold PBS, scraped off of the plates with cell scrapers, resuspended in 1mL cold PBS, and centrifuged at 300xg for 5 min at 4°C. Cell pellets were snap frozen in liquid nitrogen and stored at −80°C. RNA extraction and bulk RNA sequencing was conducted by the CHOP Center for Applied Genomics (CAG) core. In brief, RNA extraction was performed using AllPrep DNA/RNA Micro Kit (QIAGEN). RNA eluate concentration was quantified using the QUBIT HS RNA kit (Thermo Fisher) and quality was assessed by TapeStation (Agilent). All sample libraries were prepared using Illumina Stranded Total RNAseq library prep kit with 300ng of total RNA input to start. The libraries were then sequenced using a v1.5 S1 200 cycles kit on the NovaSeq 6000 at sequencing parameters 101×10×10×101 and FASTQ files were released.

The FASTQ files were subjected to quality check as described^70, 71^. The reads were aligned with Hisat2^72^ to the hg38 reference genome. The feature counts were then extracted using featureCounts^73^ followed by differential gene expression analysis using Deseq2 R package^74^. DE genes were analyzed using a hypergeometric test and the Benjamini & Hochberg method. Transformed read counts were used to make expression boxplots.

## Data availability

Raw sequencing data from patient samples (snRNA-seq, snATAC-seq) were deposited in dbGAP with accession number phs002614.v2.p1. This study did not generate any unique code. All software tools used in this study are publicly available. The authors declare that all R scripts supporting the findings of this study are available from the corresponding author upon reasonable request.

## Supporting information

Combined Supplementary Figures and Methods

## Acknowledgements

We would first and foremost like to thank the patients and families who are members of the BWS Registry and provided their tissue samples for research purposes. We thank CHOP CAG, for coordinating sequencing. We thank Dr. Kate Creasy and Amrith Rodrigues from Dr. Daniel Rader’s lab at the University of Pennsylvania for their assistance with optimizing the single-nuclei extraction from liver samples. We thank Sanam Kavari for her assistance with data upload to GEO. We also thank Dr. Rebecca Linn from the Division of Anatomic Pathology at CHOP for coordinating the nonBWS liver samples used in this study; we would also like to respectfully acknowledge the individuals from whom these samples were derived. This work was supported by NIH CA193915, a Damon Runyon Clinical Investigator Award supported by the Damon Runyon Cancer Research Foundation (105-19), Alex’s Lemonade Stand Foundation, St Baldrick’s Foundation Research Grant Award, Rally Foundation for Childhood Cancer Research Career Development Award, the Lorenzo “Turtle” Sartini, Jr. Endowed Chair in Beckwith-Wiedemann Syndrome Research, and the Victoria Fertitta Fund through the Lorenzo “Turtle” Sartini Jr. Endowed Chair in Beckwith-Wiedemann Syndrome Research; all of which were awarded to JMK. The FDA contract 75F40121C00137 to KMB.

## Author contributions

S.N., E.D.T., and J.M.K. designed the study. S.N., E.D.T., and R.D.P. performed experiments. S.N., E.D.T., R.D.P., M.B., S.M, and M.X. analyzed the data. B.M., L.G.G, and K.M.B provided materials and clinical data. S.N., E.D.T., and J.M.K. wrote the manuscript.

## References

1. Nirgude S, Naveh NSS, Kavari SL, Traxler EM, Kalish JM. Cancer predisposition signaling in Beckwith-Wiedemann Syndrome drives Wilms tumor development. Br J Cancer, (2023).

2. Klein SD, DeMarchis M, Linn RL, MacFarland SP, Kalish JM. Occurrence of Hepatoblastomas in Patients with Beckwith-Wiedemann Spectrum (BWSp). Cancers (Basel) 15, (2023).

3. Rakhshandehroo M, Knoch B, Müller M, Kersten S. Peroxisome Proliferator-Activated Receptor Alpha Target Genes. PPAR Research 2010, 612089 (2010).

4. Sobel Naveh NS, Traxler EM, Duffy KA, Kalish JM. Molecular networks of hepatoblastoma predisposition and oncogenesis in Beckwith-Wiedemann syndrome. Hepatol Commun 6, 2132–2146 (2022).

5. Duffy KA, et al. Characterization of the Beckwith-Wiedemann spectrum: Diagnosis and management. Am J Med Genet C Semin Med Genet 181, 693–708 (2019).

6. Si-Tayeb K, Lemaigre FP, Duncan SA. Organogenesis and development of the liver. Dev Cell 18, 175–189 (2010).

7. Mitra V, Metcalf J. Metabolic functions of the liver. Anaesthesia & Intensive Care Medicine 10, 334–335 (2009).

8. Rui L. Energy metabolism in the liver. Compr Physiol 4, 177–197 (2014).

9. MacParland SA, et al. Single cell RNA sequencing of human liver reveals distinct intrahepatic macrophage populations. Nat Commun 9, 4383 (2018).

10. Andrews TS, et al. Single-Cell, Single-Nucleus, and Spatial RNA Sequencing of the Human Liver Identifies Cholangiocyte and Mesenchymal Heterogeneity. Hepatol Commun 6, 821–840 (2022).

11. Wesley BT, et al. Single-cell atlas of human liver development reveals pathways directing hepatic cell fates. Nat Cell Biol 24, 1487–1498 (2022).

12. Payen VL, et al. Single-cell RNA sequencing of human liver reveals hepatic stellate cell heterogeneity. JHEP Rep 3, 100278 (2021).

13. Pilet J, et al. Preneoplastic liver colonization by 11p15.5 altered mosaic cells in young children with hepatoblastoma. Nat Commun 14, 7122 (2023).

14. Duffy KA, Cohen JL, Elci OU, Kalish JM. Development of the Serum alpha-Fetoprotein Reference Range in Patients with Beckwith-Wiedemann Spectrum. J Pediatr 212, 195–200 e192 (2019).

15. Muto Y, et al. Single cell transcriptional and chromatin accessibility profiling redefine cellular heterogeneity in the adult human kidney. Nat Commun 12, 2190 (2021).

16. Chang S, Bartolomei MS. Modeling human epigenetic disorders in mice: Beckwith-Wiedemann syndrome and Silver-Russell syndrome. Dis Model Mech 13, (2020).

17. Baker SW, Duffy KA, Richards-Yutz J, Deardorff MA, Kalish JM, Ganguly A. Improved molecular detection of mosaicism in Beckwith-Wiedemann Syndrome. J Med Genet 58, 178–184 (2021).

18. Hao Y, et al. Integrated analysis of multimodal single-cell data. Cell 184, 3573–3587 e3529 (2021).

19. Stuart T, et al. Comprehensive Integration of Single-Cell Data. Cell 177, 1888–1902 e1821 (2019).

20. Butler A, Hoffman P, Smibert P, Papalexi E, Satija R. Integrating single-cell transcriptomic data across different conditions, technologies, and species. Nat Biotechnol 36, 411–420 (2018).

21. Satija R, Farrell JA, Gennert D, Schier AF, Regev A. Spatial reconstruction of single-cell gene expression data. Nat Biotechnol 33, 495–502 (2015).

22. Korsunsky I, et al. Fast, sensitive and accurate integration of single-cell data with Harmony. Nat Methods 16, 1289–1296 (2019).

23. Panday R, Monckton CP, Khetani SR. The Role of Liver Zonation in Physiology, Regeneration, and Disease. Semin Liver Dis 42, 1–16 (2022).

24. Manco R, Itzkovitz S. Liver zonation. J Hepatol 74, 466–468 (2021).

25. Everman DB, Shuman C, Dzolganovski B, O’Riordan M A, Weksberg R, Robin NH. Serum alpha-fetoprotein levels in Beckwith-Wiedemann syndrome. J Pediatr 137, 123–127 (2000).

26. Granja JM, et al. ArchR is a scalable software package for integrative single-cell chromatin accessibility analysis. Nat Genet 53, 403–411 (2021).

27. Changizi Z, Kajbaf F, Moslehi A. An Overview of the Role of Peroxisome Proliferator-activated Receptors in Liver Diseases. J Clin Transl Hepatol 11, 1542–1552 (2023).

28. Brocker CN, Yue J, Kim D, Qu A, Bonzo JA, Gonzalez FJ. Hepatocyte-specific PPARA expression exclusively promotes agonist-induced cell proliferation without influence from nonparenchymal cells. Am J Physiol Gastrointest Liver Physiol 312, G283–G299 (2017).

29. Fan S, et al. YAP-TEAD mediates PPAR alpha-induced hepatomegaly and liver regeneration in mice. Hepatology 75, 74–88 (2022).

30. Shan J, et al. Tcf7l1 Acts as a Suppressor for the Self-Renewal of Liver Cancer Stem Cells and Is Regulated by IGF/MEK/ERK Signaling Independent of beta-Catenin. Stem Cells 37, 1389–1400 (2019).

31. Pu XY, Zheng DF, Lv T, Zhou YJ, Yang JY, Jiang L. Overexpression of transcription factor 3 drives hepatocarcinoma development by enhancing cell proliferation via activating Wnt signaling pathway. Hepatobiliary Pancreat Dis Int 21, 378–386 (2022).

32. Todisco S, et al. PPAR Alpha as a Metabolic Modulator of the Liver: Role in the Pathogenesis of Nonalcoholic Steatohepatitis (NASH). Biology (Basel) 11, (2022).

33. Aibara D, et al. Gene repression through epigenetic modulation by PPARA enhances hepatocellular proliferation. iScience 25, 104196 (2022).

34. Spangenburg EE, Pratt SJP, Wohlers LM, Lovering RM. Use of BODIPY (493/503) to visualize intramuscular lipid droplets in skeletal muscle. J Biomed Biotechnol 2011, 598358 (2011).

35. Qiu B, Simon MC. BODIPY 493/503 Staining of Neutral Lipid Droplets for Microscopy and Quantification by Flow Cytometry. Bio Protoc 6, (2016).

36. Tian G, et al. Voltage-dependent anion channel 1 (VDAC1) overexpression alleviates cardiac fibroblast activation in cardiac fibrosis via regulating fatty acid metabolism. Redox Biol 67, 102907 (2023).

37. van der Weijden VA, et al. FOXO1-mediated lipid metabolism maintains mammalian embryos in dormancy. Nat Cell Biol 26, 181–193 (2024).

38. Rosca MG, Vazquez EJ, Chen Q, Kerner J, Kern TS, Hoppel CL. Oxidation of fatty acids is the source of increased mitochondrial reactive oxygen species production in kidney cortical tubules in early diabetes. Diabetes 61, 2074–2083 (2012).

39. Chen Z, Tian R, She Z, Cai J, Li H. Role of oxidative stress in the pathogenesis of nonalcoholic fatty liver disease. Free Radical Biology and Medicine 152, 116–141 (2020).

40. Miyamoto HD, et al. Iron Overload via Heme Degradation in the Endoplasmic Reticulum Triggers Ferroptosis in Myocardial Ischemia-Reperfusion Injury. JACC Basic Transl Sci 7, 800–819 (2022).

41. Tanei T, et al. Cascade Reaction in Human Live Tissue Allows Clinically Applicable Diagnosis of Breast Cancer Morphology. Adv Sci (Weinh) 6, 1801479 (2019).

42. Wang Y, Nakajima T, Gonzalez FJ, Tanaka N. PPARs as Metabolic Regulators in the Liver: Lessons from Liver-Specific PPAR-Null Mice. Int J Mol Sci 21, (2020).

43. Annunziato S, Tchorz JS. Liver zonation—a journey through space and time. Nature Metabolism 3, 7–8 (2021).

44. Cunningham RP, Porat-Shliom N. Liver Zonation - Revisiting Old Questions With New Technologies. Front Physiol 12, 732929 (2021).

45. Halpern KB, et al. Single-cell spatial reconstruction reveals global division of labour in the mammalian liver. Nature 542, 352–356 (2017).

46. Rakhshandehroo M, Knoch B, Müller M, Kersten S. Peroxisome proliferator-activated receptor alpha target genes. PPAR Res 2010, (2010).

47. Mello T, Polvani S, Galli A. Peroxisome proliferator-activated receptor and retinoic x receptor in alcoholic liver disease. PPAR Res 2009, 748174 (2009).

48. Wang L, Liu J, Miao Z, Pan Q, Cao W. Lipid droplets and their interactions with other organelles in liver diseases. The International Journal of Biochemistry & Cell Biology 133, 105937 (2021).

49. Talari NK, et al. Lipid-droplet associated mitochondria promote fatty-acid oxidation through a distinct bioenergetic pattern in male Wistar rats. Nature Communications 14, 766 (2023).

50. Qu Q, Zeng F, Liu X, Wang QJ, Deng F. Fatty acid oxidation and carnitine palmitoyltransferase I: emerging therapeutic targets in cancer. Cell Death Dis 7, e2226 (2016).

51. Hayes JD, Dinkova-Kostova AT, Tew KD. Oxidative Stress in Cancer. Cancer Cell 38, 167–197 (2020).

52. Barrera G. Oxidative stress and lipid peroxidation products in cancer progression and therapy. ISRN Oncol 2012, 137289 (2012).

53. Harris IS, DeNicola GM. The Complex Interplay between Antioxidants and ROS in Cancer. Trends Cell Biol 30, 440–451 (2020).

54. Wang R, Hao W, Pan L, Boldogh I, Ba X. The roles of base excision repair enzyme OGG1 in gene expression. Cell Mol Life Sci 75, 3741–3750 (2018).

55. Su LJ, et al. Reactive Oxygen Species-Induced Lipid Peroxidation in Apoptosis, Autophagy, and Ferroptosis. Oxid Med Cell Longev 2019, 5080843 (2019).

56. Juric-Sekhar G, Zarkovic K, Waeg G, Cipak A, Zarkovic N. Distribution of 4-hydroxynonenal-protein conjugates as a marker of lipid peroxidation and parameter of malignancy in astrocytic and ependymal tumors of the brain. Tumori 95, 762–768 (2009).

57. Skrzydlewska E, Stankiewicz A, Sulkowska M, Sulkowski S, Kasacka I. Antioxidant status and lipid peroxidation in colorectal cancer. J Toxicol Environ Health A 64, 213–222 (2001).

58. Oberley TD, Toyokuni S, Szweda LI. Localization of hydroxynonenal protein adducts in normal human kidney and selected human kidney cancers. Free Radical Biology and Medicine 27, 695–703 (1999).

59. Karihtala P, Kauppila S, Puistola U, Jukkola-Vuorinen A. Divergent behaviour of oxidative stress markers 8-hydroxydeoxyguanosine (8-OHdG) and 4-hydroxy-2-nonenal (HNE) in breast carcinogenesis. Histopathology 58, 854–862 (2011).

60. Allameh A, Niayesh-Mehr R, Aliarab A, Sebastiani G, Pantopoulos K. Oxidative Stress in Liver Pathophysiology and Disease. Antioxidants (Basel) 12, (2023).

61. Gandhi CR. Oxidative Stress and Hepatic Stellate Cells: A PARADOXICAL RELATIONSHIP. Trends Cell Mol Biol 7, 1–10 (2012).

62. Tan Y, Wang M, Yang K, Chi T, Liao Z, Wei P. PPAR-alpha Modulators as Current and Potential Cancer Treatments. Front Oncol 11, 599995 (2021).

63. McGinnis CS, Murrow LM, Gartner ZJ. DoubletFinder: Doublet Detection in Single-Cell RNA Sequencing Data Using Artificial Nearest Neighbors. Cell Syst 8, 329–337 e324 (2019).

64. Aran D, et al. Reference-based analysis of lung single-cell sequencing reveals a transitional profibrotic macrophage. Nat Immunol 20, 163–172 (2019).

65. Ianevski A, Giri AK, Aittokallio T. Fully-automated and ultra-fast cell-type identification using specific marker combinations from single-cell transcriptomic data. Nat Commun 13, 1246 (2022).

66. Bibby JA, et al. Systematic single-cell pathway analysis to characterize early T cell activation. Cell Rep 41, 111697 (2022).

67. Bravo Gonzalez-Blas C, et al. SCENIC+: single-cell multiomic inference of enhancers and gene regulatory networks. Nat Methods 20, 1355–1367 (2023).

68. Shannon P, et al. Cytoscape: a software environment for integrated models of biomolecular interaction networks. Genome Res 13, 2498–2504 (2003).

69. Yang W, et al. Generation of iPSCs as a Pooled Culture Using Magnetic Activated Cell Sorting of Newly Reprogrammed Cells. PLoS One 10, e0134995 (2015).

70. Nirgude S, Desai S, Choudhary B. Curcumin alters distinct molecular pathways in breast cancer subtypes revealed by integrated miRNA/mRNA expression analysis. Cancer Rep (Hoboken) 5, e1596 (2022).

71. Nirgude S, Desai S, Mahadeva R, Ravindran F, Choudhary B. ST08 Altered NF-kappaB Pathway in Breast Cancer Cells In Vitro as Revealed by miRNA-mRNA Analysis and Enhanced the Effect of Cisplatin on Tumour Reduction in EAC Mouse Model. Front Oncol 12, 835027 (2022).

72. Kim D, Paggi JM, Park C, Bennett C, Salzberg SL. Graph-based genome alignment and genotyping with HISAT2 and HISAT-genotype. Nat Biotechnol 37, 907–915 (2019).

73. Liao Y, Smyth GK, Shi W. featureCounts: an efficient general purpose program for assigning sequence reads to genomic features. Bioinformatics 30, 923–930 (2014).

74. Love MI, Huber W, Anders S. Moderated estimation of fold change and dispersion for RNA-seq data with DESeq2. Genome Biol 15, 550 (2014).

