## Supplementary material for "Single-nucleus multiomic analysis of Beckwith-Wiedemann syndrome liver reveals PPARA signaling enrichment and metabolic dysfunction": Combined Supplementary Figures and Methods

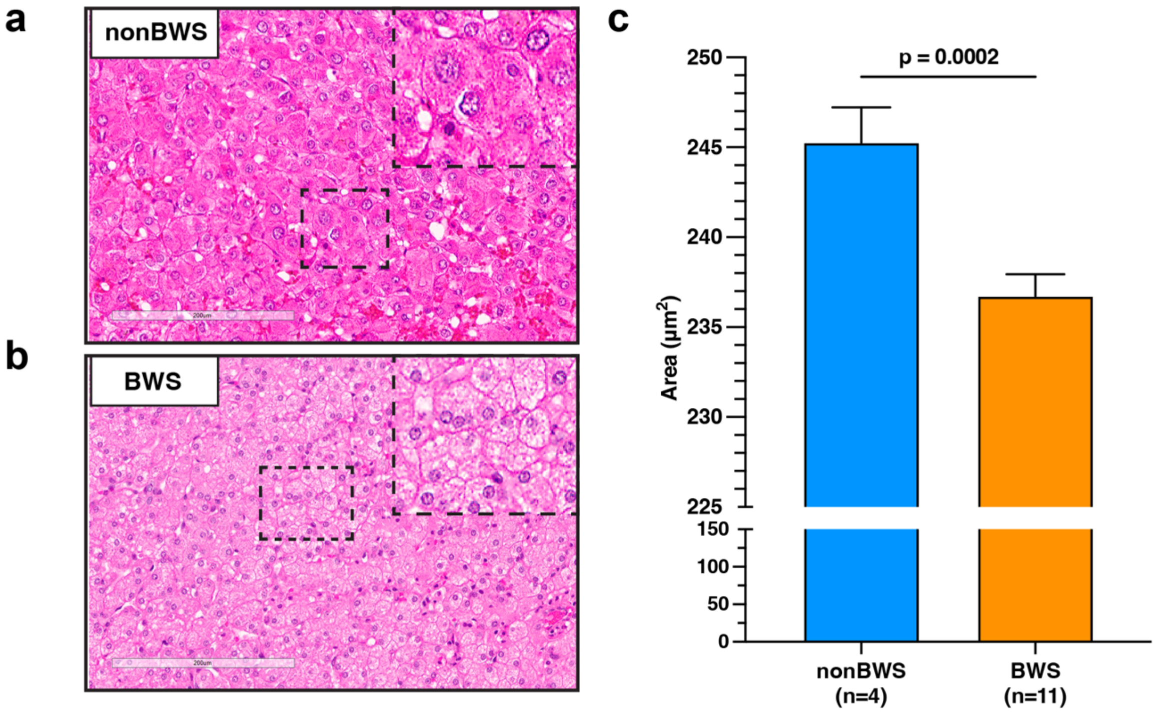


**Supplementary Figure 1: Assessment of nonBWS and BWS normal liver histology.** Representative liver sections from nonBWS (**a**; control normal liver) and BWS (**b**; normal liver) were stained with hematoxylin and eosin. Scale bar represents 200 µm. Hepatocyte size was determined from 4 control and 11 BWS livers. Displayed are the mean ± SEM. Data were analyzed with an unpaired student t-test.

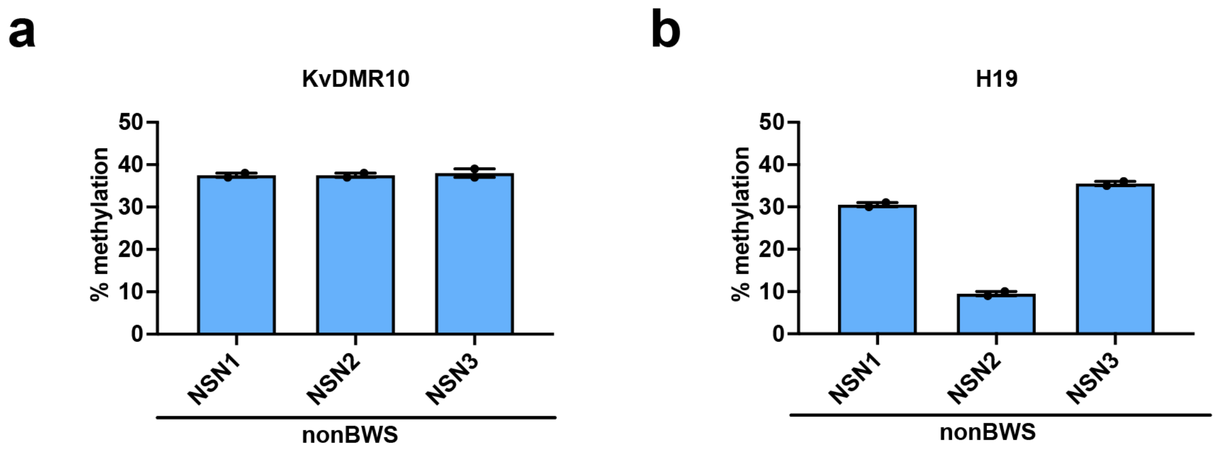


**Supplementary Figure 2**: Methylation values of IC2 (KvDMR10; **a**) and IC1 (H19; **b**), measured by bisulfite pyrosequencing the bisulfite converted DNA from liver tissues in study. Displayed are mean ± SEM.

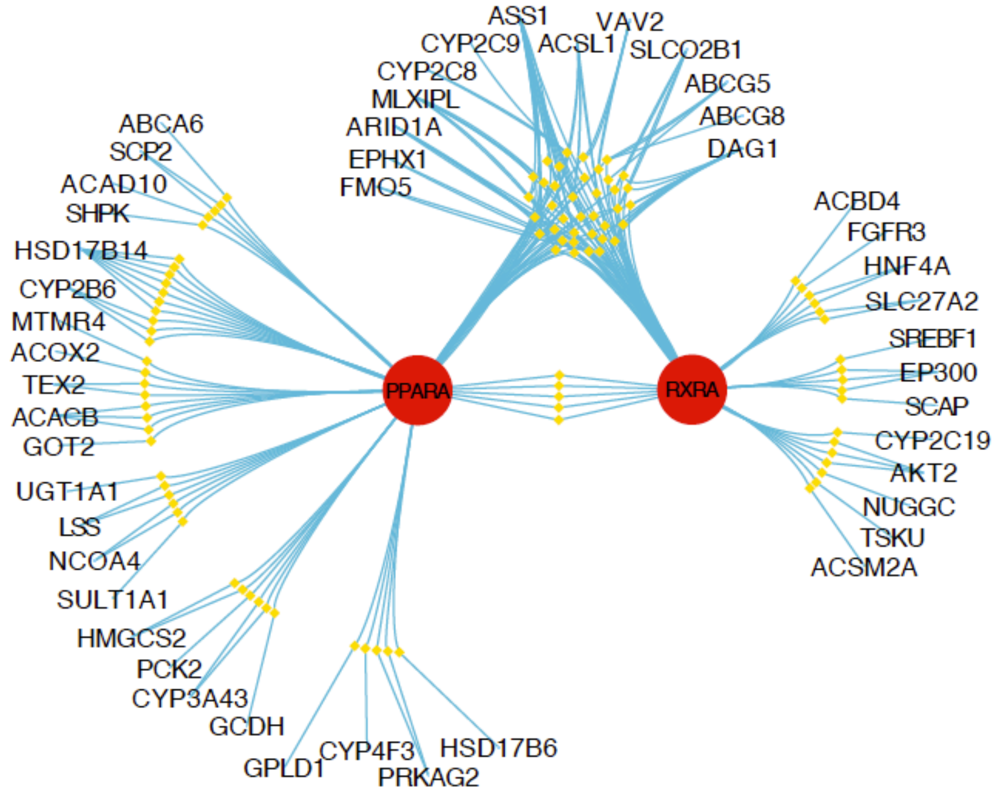


**Supplementary Figure 3:** Interaction graph of transcription factor – putative enhancer – gene from PPARA and RXRA eRegulons by comparing hepatocytes (cluster 0, 1, 7) of BWS and nonBWS cohorts. Only genes are associated with the top 8000 most variable accessible regions that fall under GO terms – Cellular response to lipid, lipid localization, fatty acid metabolic process, Regulation of lipid metabolic process and steroid metabolic process are plotted. In this network, genomic regions are represented by diamonds, and the interaction between elements is represented by lines.

**a**

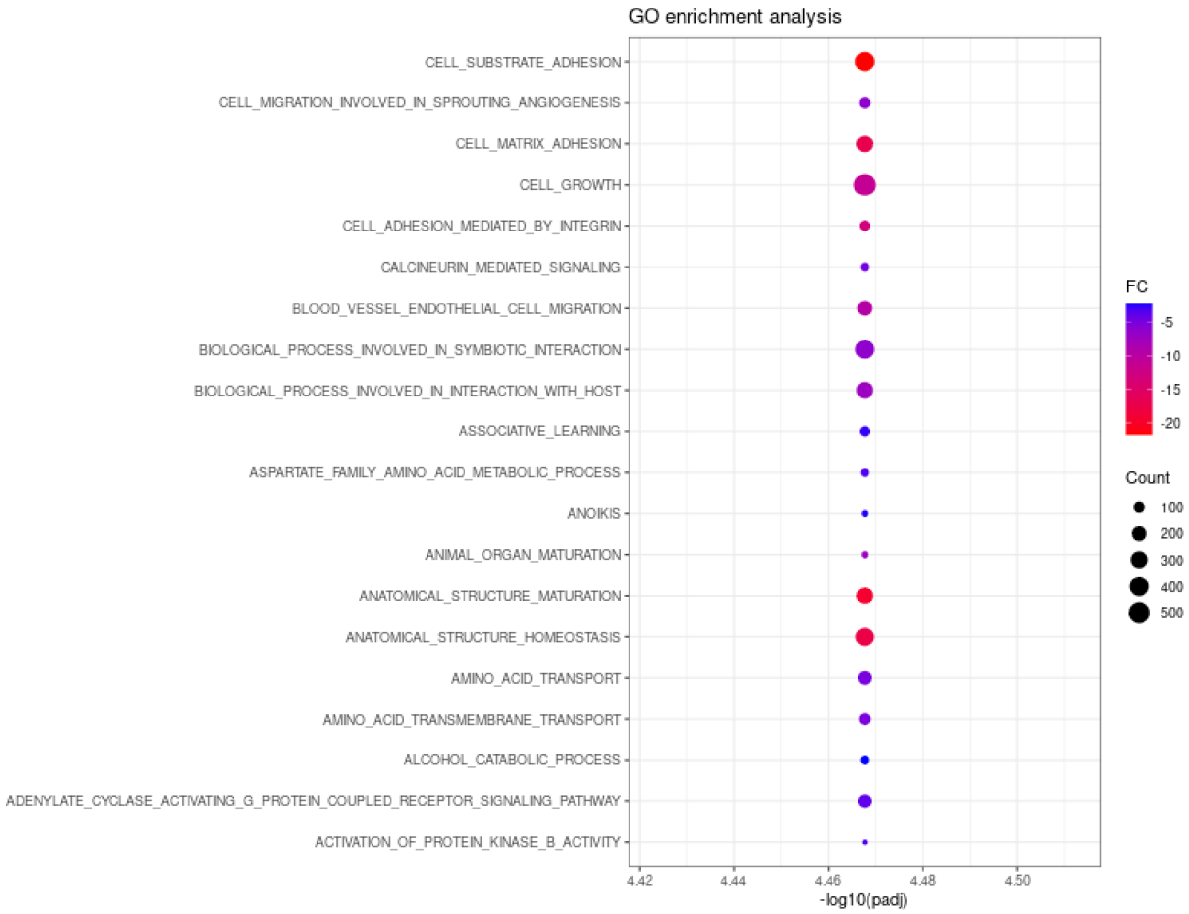

GO enrichment analysis
CELL_SUBSTRATE_ADHESION
CELL_MIGRATION_INVOLVED_IN_SPROUTING_ANGIOGENESIS
CELL_MATRIX_ADHESION
CELL_GROWTH
CELL_ADHESION_MEDIATED_BY_INTEGRIN
CALCINEURIN_MEDIATED_SIGNALING
BLOOD_VESSEL_ENDOTHELIAL_CELL_MIGRATION
BIOLOGICAL_PROCESS_INVOLVED_IN_SYMBIOTIC_INTERACTION
BIOLOGICAL_PROCESS_INVOLVED_IN_INTERACTION_WITH_HOST
ASSOCIATIVE_LEARNING
ASPARTATE_FAMILY_AMINO_ACID_METABOLIC_PROCESS
ANOIKIS
ANIMAL_ORGAN_MATURATION
ANATOMICAL_STRUCTURE_MATURATION
ANATOMICAL_STRUCTURE_HOMEOSTASIS
AMINO_ACID_TRANSPORT
AMINO_ACID_TRANSMEMBRANE_TRANSPORT
ALCOHOL_CATABOLIC_PROCESS
ADENYLATE_CYCLASE_ACTIVATING_G_PROTEIN_COUPLED_RECEPTOR_SIGNALING_PATHWAY
ACTIVATION_OF_PROTEIN_KINASE_B_ACTIVITY
4.42
4.44
4.46
4.48
4.50
-log10(padj)
FC
-5
-10
-15
-20
Count
100
200
300
400
500

**b**

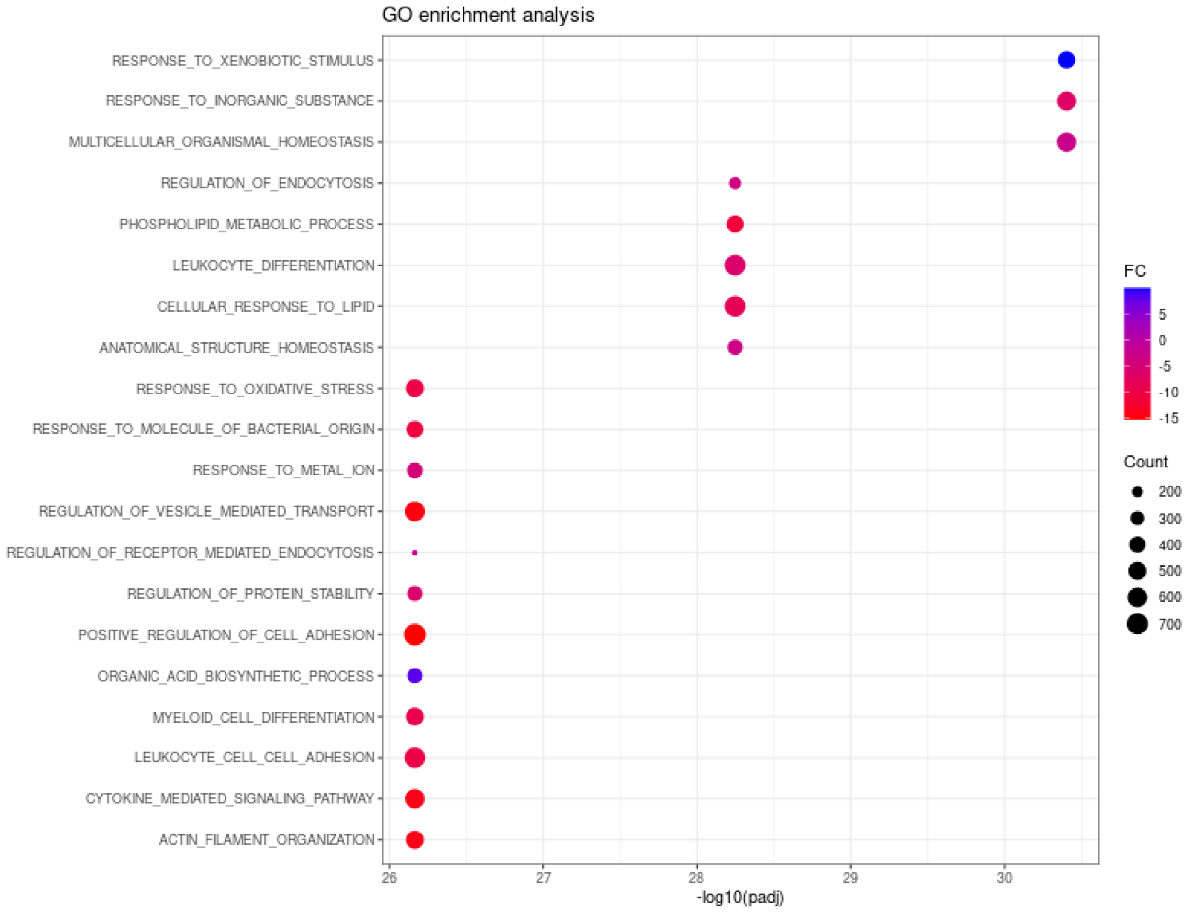

GO enrichment analysis
RESPONSE_TO_XENOBIOTIC_STIMULUS
RESPONSE_TO_INORGANIC_SUBSTANCE
MULTICELLULAR_ORGANISMAL_HOMEOSTASIS
REGULATION_OF_ENDOCYTOSIS
PHOSPHOLIPID_METABOLIC_PROCESS
LEUKOCYTE_DIFFERENTIATION
CELLULAR_RESPONSE_TO_LIPID
ANATOMICAL_STRUCTURE_HOMEOSTASIS
RESPONSE_TO_OXIDATIVE_STRESS
RESPONSE_TO_MOLECULE_OF_BACTERIAL_ORIGIN
RESPONSE_TO_METAL_ION
REGULATION_OF_VESICLE_MEDIATED_TRANSPORT
REGULATION_OF_RECEPTOR_MEDIATED_ENDOCYTOSIS
REGULATION_OF_PROTEIN_STABILITY
POSITIVE_REGULATION_OF_CELL_ADHESION
ORGANIC_ACID_BIOSYNTHETIC_PROCESS
MYELOID_CELL_DIFFERENTIATION
LEUKOCYTE_CELL_CELL_ADHESION
CYTOKINE_MEDIATED_SIGNALING_PATHWAY
ACTIN_FILAMENT_ORGANIZATION
26
27
28
29
30
-log10(padj)
FC
5
0
-5
-10
-15
Count
200
300
400
500
600
700

c

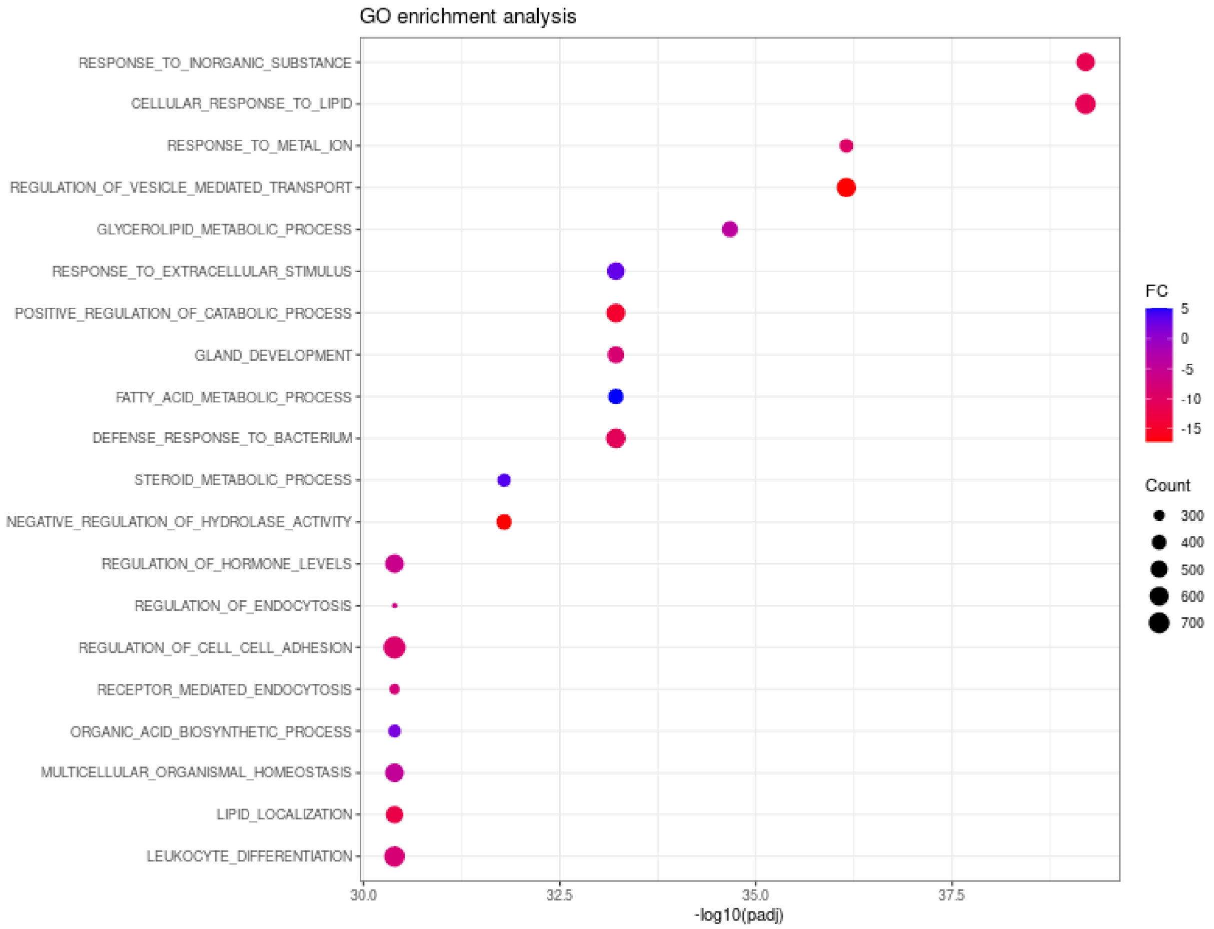
GO enrichment analysis
RESPONSE_TO_INORGANIC_SUBSTANCE
CELLULAR_RESPONSE_TO_LIPID
RESPONSE_TO_METAL_ION
REGULATION_OF_VESICLE_MEDIATED_TRANSPORT
GLYCEROLIPID_METABOLIC_PROCESS
RESPONSE_TO_EXTRACELLULAR_STIMULUS
POSITIVE_REGULATION_OF_CATABOLIC_PROCESS
GLAND_DEVELOPMENT
FATTY_ACID_METABOLIC_PROCESS
DEFENSE_RESPONSE_TO_BACTERIUM
STEROID_METABOLIC_PROCESS
NEGATIVE_REGULATION_OF_HYDROLASE_ACTIVITY
REGULATION_OF_HORMONE_LEVELS
REGULATION_OF_ENDOCYTOSIS
REGULATION_OF_CELL_CELL_ADHESION
RECEPTOR_MEDIATED_ENDOCYTOSIS
ORGANIC_ACID_BIOSYNTHETIC_PROCESS
MULTICELLULAR_ORGANISMAL_HOMEOSTASIS
LIPID_LOCALIZATION
LEUKOCYTE_DIFFERENTIATION
30.0
32.5
35.0
37.5
-log10(padj)
FC
5
0
-5
-10
-15
Count
300
400
500
600
700

**d**

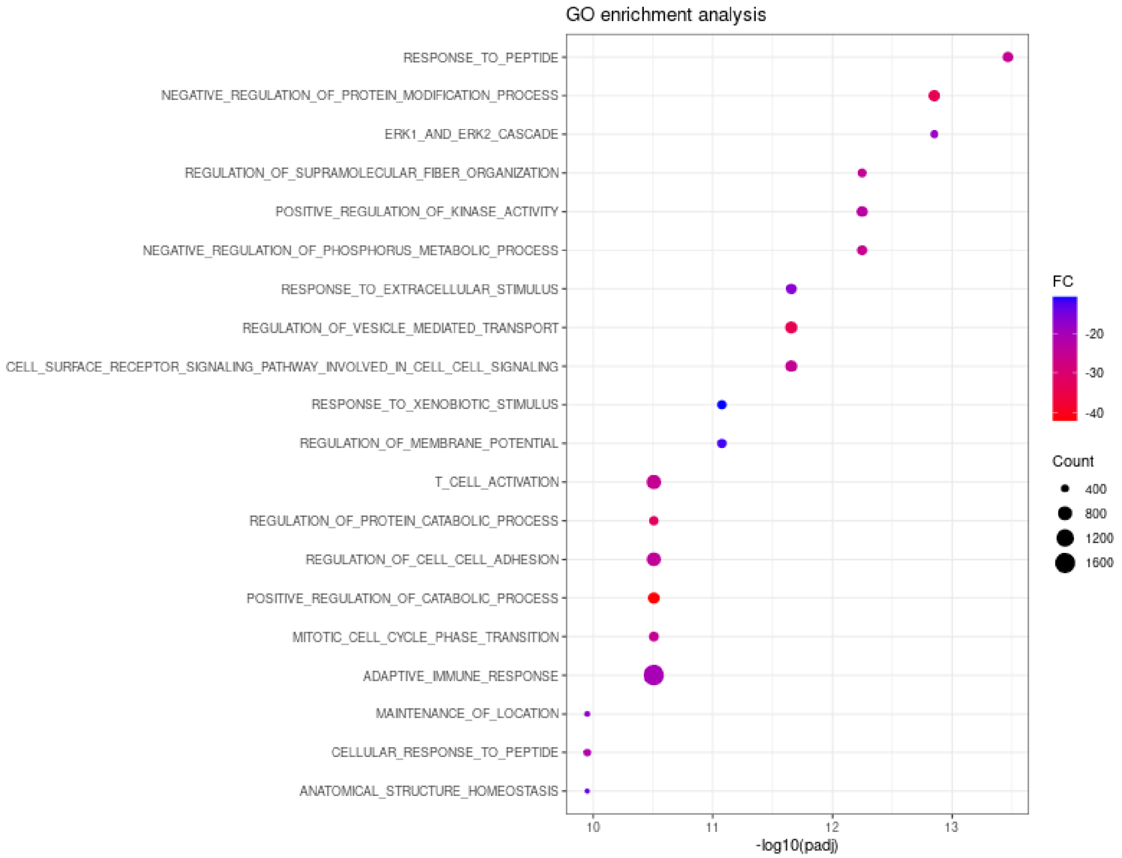


**Supplementary Figure 4:** GO terms enriched in the hepatocytes clusters (Cluster 17, 14, 15, 12) (**a-d**) of BWS liver cohort when compared to nonBWS liver cohort.

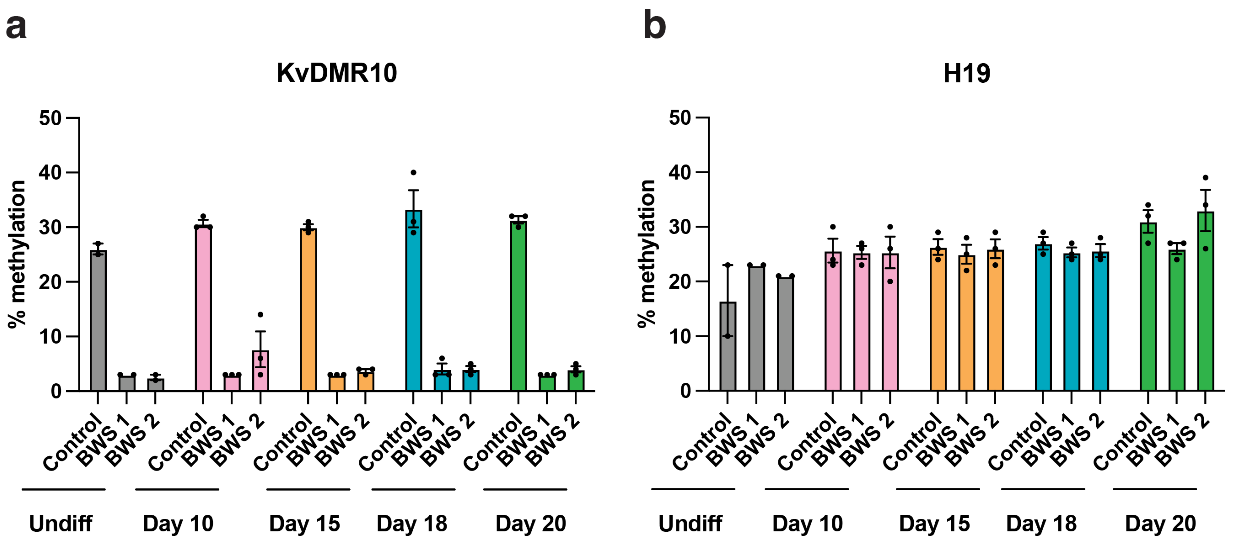


**Supplementary Figure 5:** Methylation values of IC2 (KvDMR10; **a**) and IC1 (H19; **b**), measured by bisulfite pyrosequencing the bisulfite converted DNA from iPSCs collected at different stages of hepatocyte differentiation. Displayed are mean ± SEM.

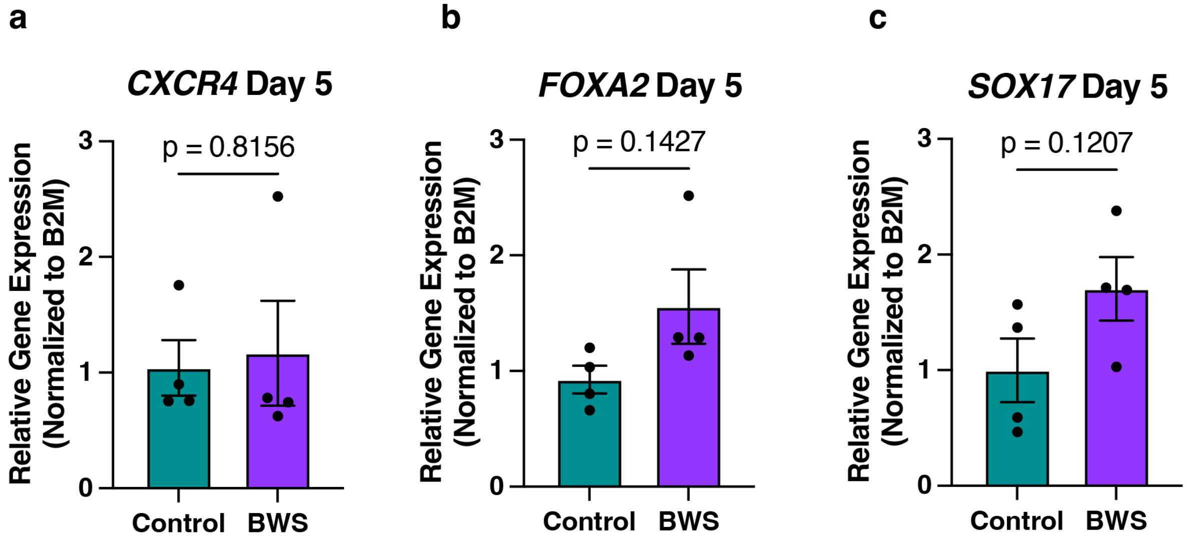


**Supplementary Figure 6: Expression of definitive endoderm markers**. Control and IC2 LOM BWS iPSCs were differentiated for 5 days in the STEMdiff hepatocyte kit. QRT-PCR of gene expression of definitive endoderm markers CXCR4 (**a**), FOXA2 (**b**), and SOX17 (**c**) were determined and normalized to expression of the B2M housekeeping gene. Each data point is the average of at least 3 technical replicates and represents one biological replicate. Displayed are mean ± S.E.M. Statistics were completed using a unpaired t-test with Welch's correction.

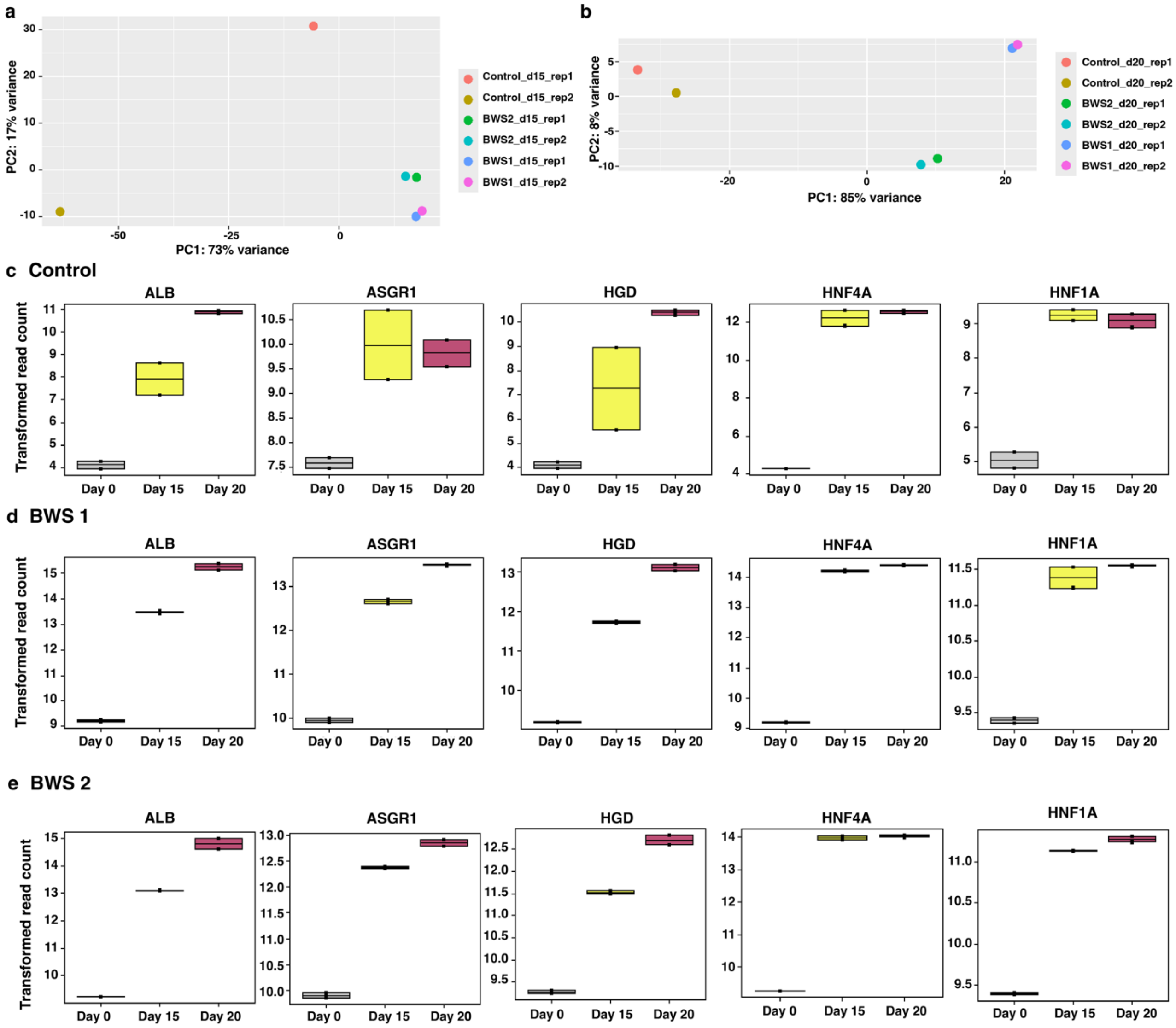


**Supplementary Figure 7: Expression of hepatocyte-specific genes from differentiated iPSCs. a&b)** PCA plots showing the variance between experimental samples from day 15 and day 20 of differentiation into hepatocytes, respectively. Samples were sequenced as biological duplicates. **c-e)** expression of hepatocyte genes albumin, asialoglycoprotein receptor, homogentisate oxidase, hepatocyte nuclear factor 4, and hepatocyte nuclear factor 1a during iPSC to hepatocyte differentiation in control parental line **(c)**, or BWS lines 1 and 2 **(d, e)**, respectively.

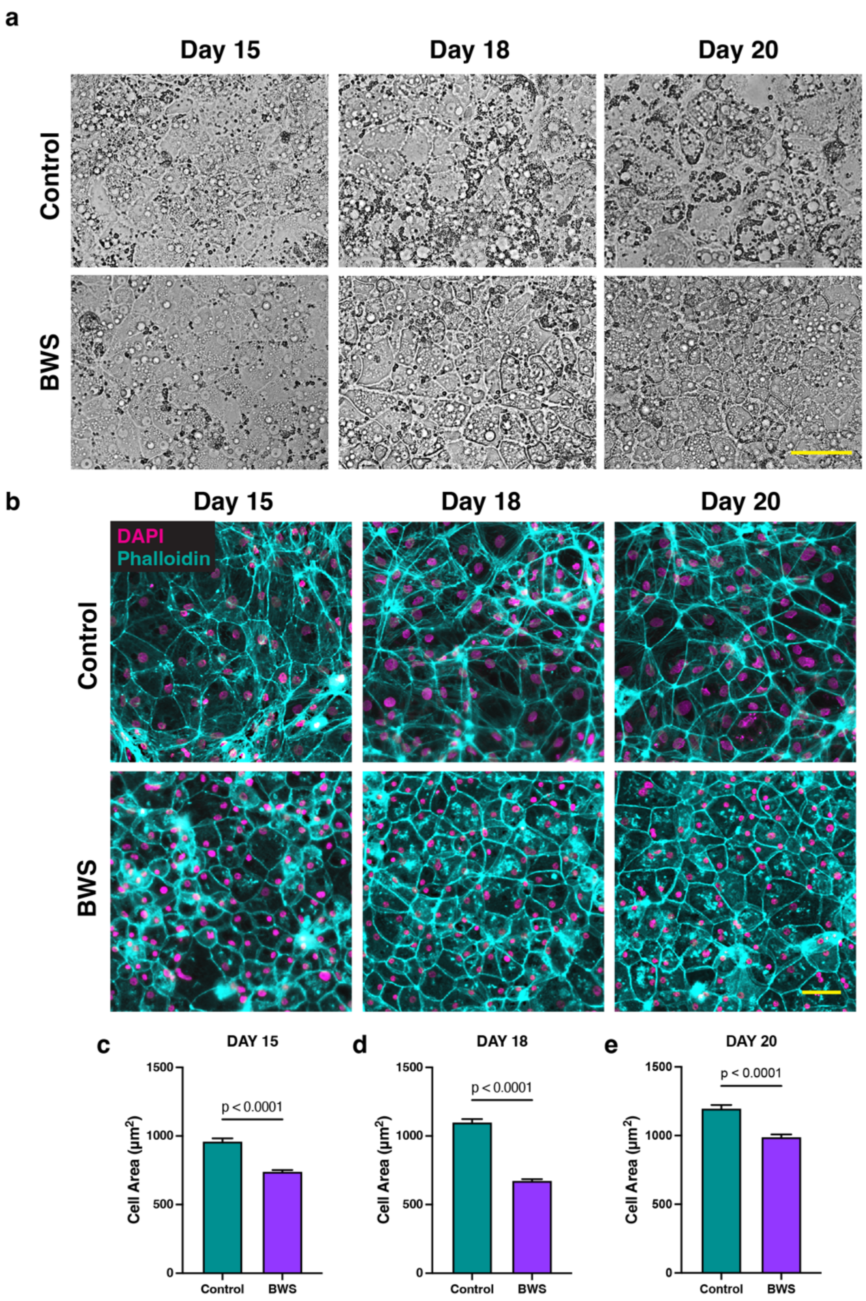


**Supplementary Figure 8: Characteristics of differentiating iPSC-derived hepatocytes. (a)** Phase contrast images of control and BWS hepatocytes at days 15, 18, and 20 of our differentiation protocol. Scale bar: 75µm**. (b)** Control and BWS hepatocytes were stained with phalloidin to stain the actin cytoskeleton at days 15, 18, and 20 of our differentiation protocol. Scale bar: 50µm. (**c-e)** Analysis of iPSC-derived hepatocyte sizes, based on phalloidin cell outlines. Displayed are mean ± SEM. p values were generated from unpaired t tests.

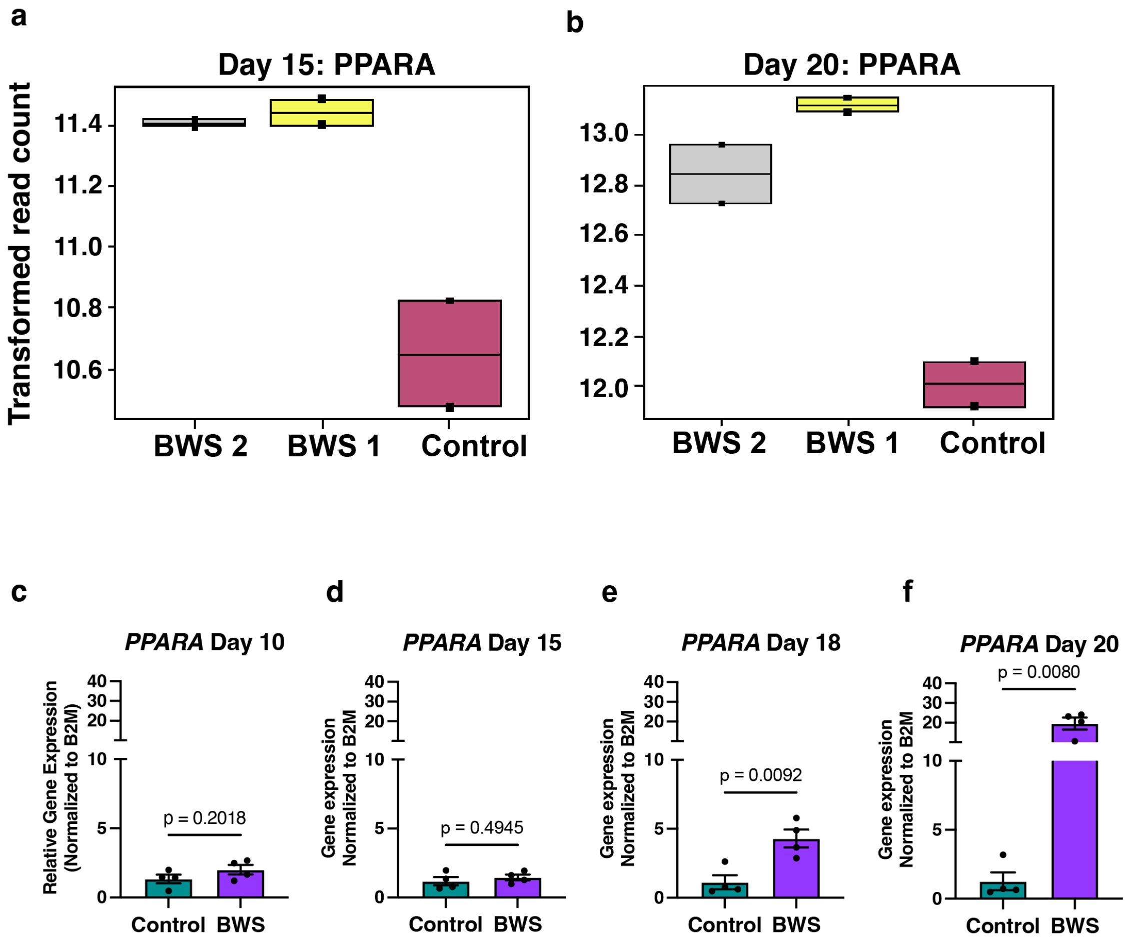


**Supplementary Figure 9: Expression of *PPARA* in iPSCs differentiated to hepatocytes in control and BWS lines. a&b)** RNAseq analysis of PPARA expression at day 15 and day 20 of our differentiation protocol, respectively. **(c-f)** Multiplexed quantitative real-time PCR of PPARA expression, relative to the B2M housekeeping gene in control and BWS 1 iPSC cells differentiated into hepatocytes and collected at different times post-differentiation induction. Each data point represents a biological replicate. Displayed are mean ± SEM, and significance was detected by unpaired t test.

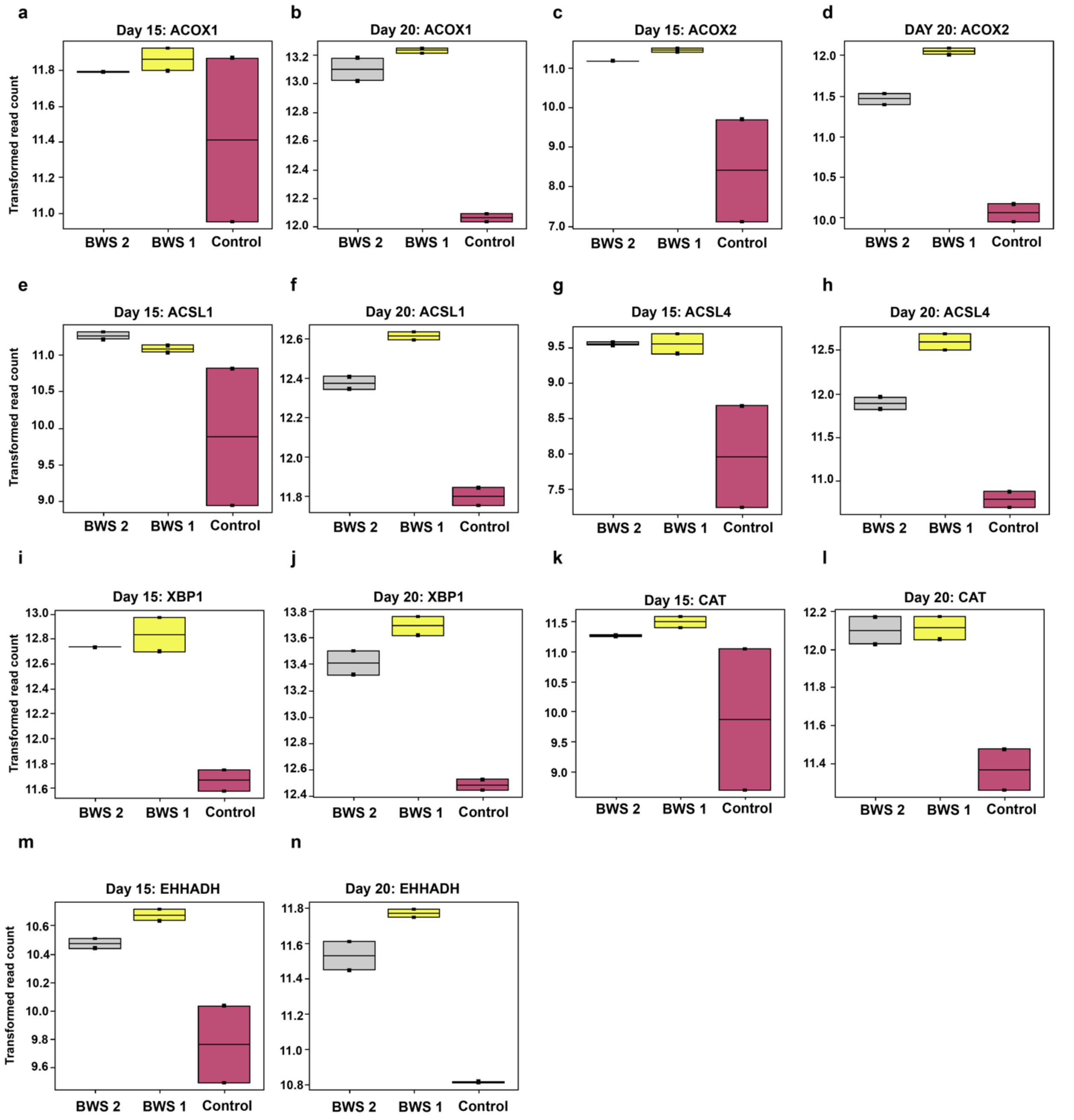


**Supplementary Figure 10: Expression of genes involved in fatty acid β-oxidation in iPSCs differentiated into hepatocytes and examined on day 15 and day 20.** We found upregulation of transcripts in BWS lines by RNAseq coding for: Acyl-CoA Oxidase 1 (**a, b**), Acyl-CoA Oxidase 2 (**c,d**), Acyl-CoA Synthetase Long Chain Family Member 1 (**e, f**), Acyl-CoA Synthetase Long Chain Family Member 4 (**g, h**), X-Box Binding Protein 1 (**i, j**) , Catalase (**k, l**), and Enoyl-CoA Hydratase and 3-Hydroxyacyl CoA Dehydrogenase (**m, n**).

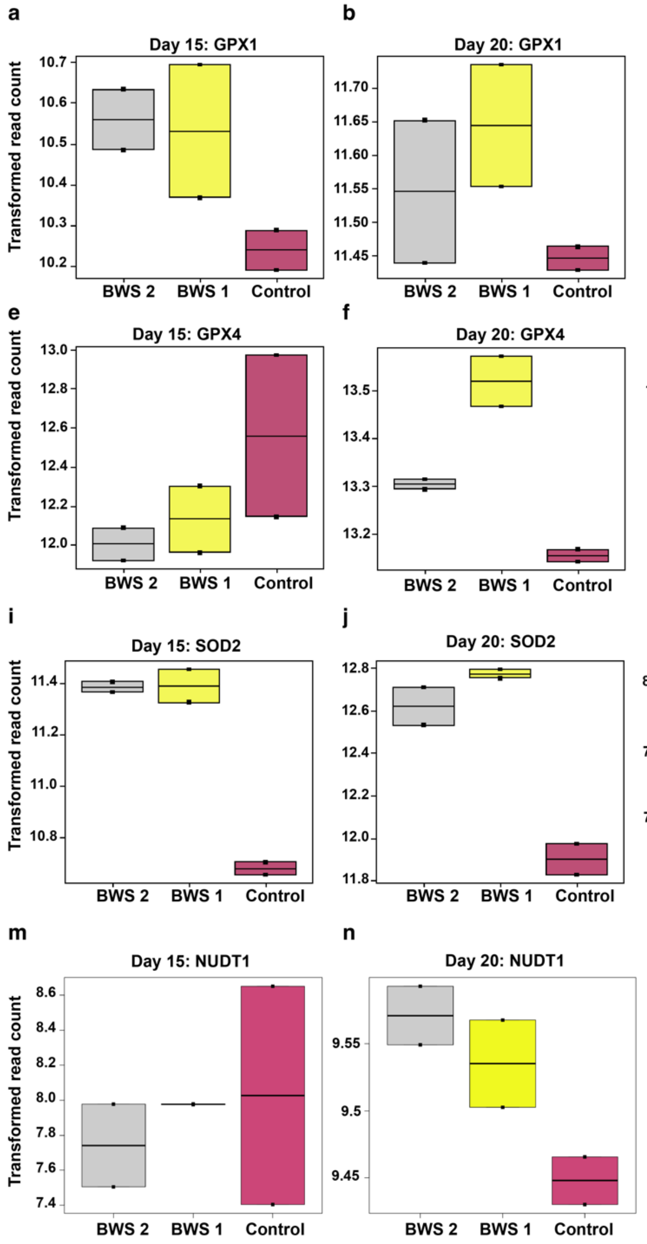


**Supplementary Figure 11: Expression of ROS-responding genes at day 15 and day 20 of iPSC to hepatocyte differentiation.** iPSCs were differentiated into hepatocytes and examined by RNAseq for expression of glutathione peroxidase 1 **(a, b)**, glutathione peroxidase 8 **(c, d)**, glutathione peroxidase 4 **(e, f)**, glutathione peroxidase 7 **(g, h)**, superoxide dismutase 2 **(i, j)**, superoxide dismutase 3 **(k, l)**, and nudix hydrolase 1 **(m, n)**.

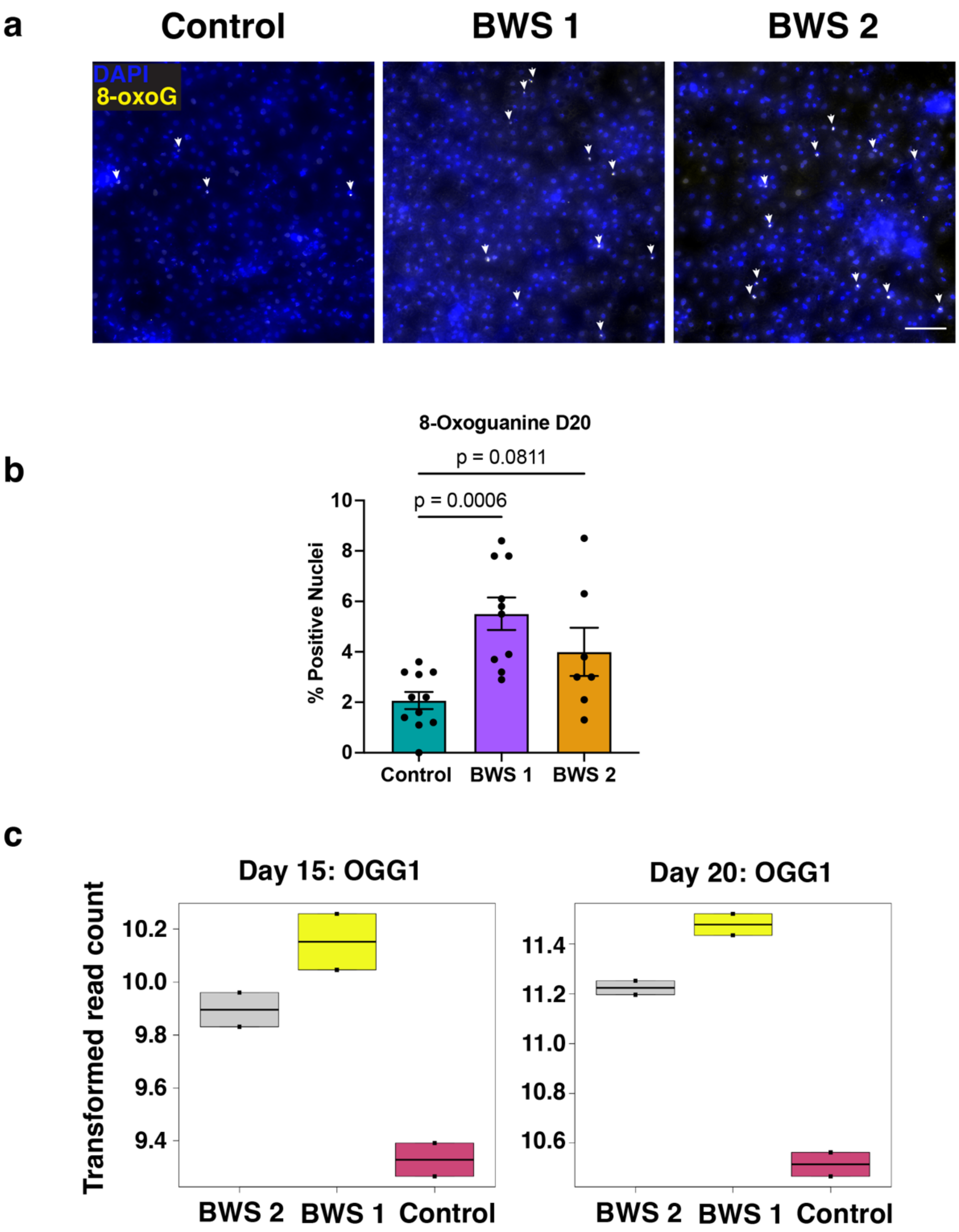


**Supplementary Figure 12: Oxidative DNA damage in BWS hepatocytes. (a)** Representative image of 8-oxoguanine staining of Day 20 iPSC to hepatocyte differentiated cells from control, BWS 1, and BWS 2 lines. Arrows demonstrate positive staining. Scale bar: 100µm. (b) Analysis of 8-oxoguanine damage from (a). Each data point represents one image. Analysis was done with at least 3 biological replicates per line. Displayed are mean ± SEM. p values were generated by one way ANOVA. (c) Expression level of 8-oxoguanine DNA glycosylase 1 transcripts in control and BWS lines at day 15 and day 20 iPSC differentiation into hepatocytes. Data are from RNA sequencing and come from 2 biological replicates.

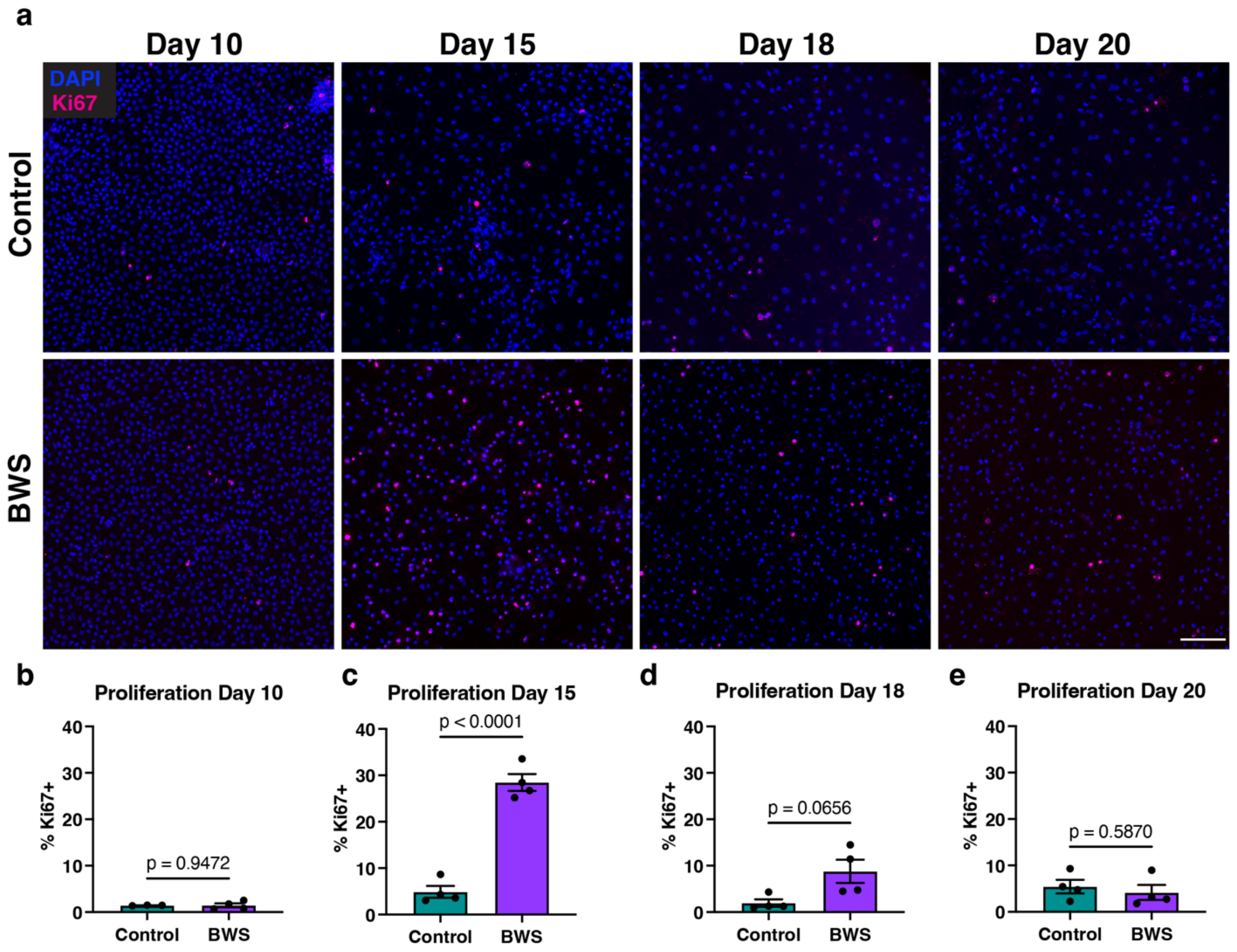


**Supplementary Figure 13: Proliferative indices during hepatocyte differentiation. (a)** Representative images of staining for Ki67 during iPSC to hepatocyte differentiation at the indicated time points. Scale bar: 50µm. (**b -d)** Quantification of Ki67 positivity at Day 10, Day 15, Day 18, and Day 20, respectively. Each data point is a biological replicate. Displayed are mean ± SEM. p values were calculated from unpaired t tests.

**Supplementary Table 1**: Clinical information on the cohort used in study.

| Sample ID | Cohort | BWS subtype in blood | Sex | Age at Surgery |
|---|---|---|---|---|
| BWSN1 | BWS | BWS IC2 LOM | F | 5 months |
| BWSN2 | BWS | BWS pUPD11 | M | 13 months |
| BWSN3 | BWS | BWS pUPD11 | F | 14 months |
| BWSN4 | BWS | BWS IC2 LOM | M | 2 months |
| NSN1 | nonBWS | nonBWS | M | 2 months |
| NSN2 | nonBWS | nonBWS | F | 1 month |
| NSN3 | nonBWS | nonBWS | M | 8 years |

**Supplementary Table 2:** Primer sequences used for pyrosequencig

| Primer's name | Sequences |
|---|---|
| H19 Forward | 5'-TGGGTATTTTTGGAGGTTTTTTT-3' |
| H19 Reverse | 5'-Biotin-TCCCATAAATATCCTATTCCCAAA-3' |
| H19 Sequencing primer | 5'-GTAGGTTTATATATTATAG-3' |
| KvDMR 10 Forward | 5'-GTATTGTTTAGGTTAGGTTGTATTG-3' |
| bio KvDMR 10 Reverse | 5'-Biotin-ACCCTCCCC ATCTCTCTA-3' |
| KVDMR 10 Sequencing primer | 5'-GGGGTATATAGTTTATTTTAGTAA-3' |

### Supplementary Methods

#### DNA extraction and Bisulfite Conversion

Genomic DNA from liver samples was isolated using the AllPrep DNA/RNA Micro Kit (Qiagen) and quantified on Qubit with HS DNA kit (Thermo Fisher). 100-500ng of DNA was bisulfite converted using the EpiTect bisulfite kit (Qiagen), according to the manufacturer’s instructions.

#### Pyrosequencing

Pyrosequencing was performed on bisulfite-converted DNA using the Pyromark PCR kit (Qiagen), and sequencing was performed on a Pyromark Q48 Autoprep (Qiagen). Sequencing primers are listed in Supplementary Table 2.

#### Histology of livers

Control (nonBWS/NSN) nontumor livers and BWS nontumor livers were FFPE processed, sectioned, and stained using standard hematoxylin and eosin staining protocols. Sections were imaged using Leica Aperio AT2 slide scanner. Hepatocyte size was manually enumerated in FIJI by investigators blinded to cohort information. At least 2 images were analyzed per patient sample.

#### Immunofluorescence staining of differentiating iPSCs

Cells were grown on coverglass and were fixed in 4% paraformaldehyde in PBS for 10 minutes and washed twice with PBS. Cells were permeabilized with 0.5% triton X-100 in PBS for 5 minutes, washed once with PBS, and blocked for 1 hour with blocking buffer (3% BSA, 0.1% triton X-100, all in PBS) at room temperature. Cells were stained with anti-Ki67 antibody (Cell Signaling #9448; 1/200) in blocking buffer overnight at 4ºC. The following day, cells were washed three times with PBS and incubated with secondary antibody (Donkey anti-mouse IgG Alexa Fluor 488;

1/1000, in blocking buffer, Fisher Scientific) for one hour at room temperature in the dark. Cells were washed three times with PBS and mounted onto slides using Prolong gold with DAPI (Fisher Scientific). For phalloidin staining, fixed cells were permeabilized with saponin (0.5% in PBS) for 10 minutes, washed with PBS, and stained with Alexa Fluor 647 Phalloidin (1/50; Cell Signaling) in PBS in the dark for 30 minutes. Cells were washed three times with PBS and mounted on slides with Prolong gold with DAPI. Cells were imaged on a Nikon Ti-2 inverted microscope. At least 3 biological replicates were analyzed per sample.

**8-oxyoguanine staining of differentiated iPSCs**

Day 20-differentiated hepatocytes were fixed in 4% PFA/PBS for 10 min, washed twice with PBS, 3 min each, and permeabilized with 0.5% triton X-100/PBS for 10 min. Cells were washed and incubated with DNase-free RNAse A (100µg/mL; Fermentas/Fisher Scientific) for 20 min at 37ºC to remove any oxidative RNA damage. Cells were washed with PBS twice, for 3 min each and then incubated with 2N HCl for 30 min at 37ºC. Acid solution was aspirated and neutralized by washing with 0.1M Borate buffer, pH 8.5 (Thermo Fisher Scientific) three times for 3 min each. Cells were subsequently washed with PBS twice for 3 min each and blocked at room temperature with 3% BSA/PBS containing 0.1% triton X-100 for one hour. Primary antibody to 8-oxyoguanine was diluted in blocking buffer (Rockland; 1/200) overnight at 4ºC. Cells were washed twice with PBS and incubated with Donkey anti-mouse Alexa Fluor 488 IgG (1/2,000) for one hour at room temperature. Cells were stained with DAPI (10µg/mL) for 5 min and washed with PBS before imaging on a Nikon Ti-2 inverted microscope. At least 3 biological replicates were analyzed per line.
